# Chronic wireless neural population recordings with common marmosets

**DOI:** 10.1101/2021.02.25.432889

**Authors:** Jeffrey D. Walker, Friederice Pirschel, Marina Sundiang, Marek Niekrasz, Jason N. MacLean, Nicholas G. Hatsopoulos

**Affiliations:** Committee on Computational Neuroscience, University of Chicago, Chicago, Illinois; Department of Organismal Biology and Anatomy, University of Chicago, Chicago, Illinois; Department of Surgery, University of Chicago, Chicago, Illinois; Department of Neurobiology, University of Chicago, Chicago, Illinois; Grossman Institute for Neuroscience, Quantitative Biology and Human Behavior, University of Chicago, Chicago, Illinois

**Keywords:** Marmoset, wireless neural recording, multi-electrode arrays, motor control, natural behavior

## Abstract

Marmosets are an increasingly important model system for neuroscience in part due to genetic tractability and enhanced cortical accessibility, due to a lissencephalic neocortex. However, many of the techniques generally employed to record neural activity in primates inhibit the expression of natural behaviors in marmosets precluding neurophysiological insights. To address this challenge, we developed methods for recording neural population activity in unrestrained marmosets across multiple ethological behaviors, multiple brain states, and over multiple years. Notably, our flexible methodological design allows for replacing electrode arrays and removal of implants providing alternative experimental endpoints. We validate the method by recording sensorimotor cortical population activity in freely moving marmosets across their natural behavioral repertoire and during sleep.

**HIGHLIGHTS:**

– Simultaneous and chronic wireless neural population recordings in multiple freely moving marmosets
– Neural recording approach enables studies of natural repertoire of behaviors and sleep
– Methyl-methacrylate free surgical approach designed to promote biocompatibility and longitudinal success of the implant
– Modular headstage configuration requires minimal daily animal handling for daily neural recordings
– Alternative experimental endpoints: implant removal, healing, and electrode array replacement

## INTRODUCTION

Marmosets show great potential as a model species for studying many aspects of nervous system function (Miller, 2017). Their fecundity and demonstrated capacity for genetic manipulation suggest they will become a key non-human primate transgenic model for circuit dissection in multiple fields of neuroscience (Belmonte et al., 2015; Sasaki, 2015; Sasaki et al., 2009). Early developments of single unit electrophysiology with marmosets led to insights about auditory processing (Eliades and Wang, 2008a, 2008b; Lu et al., 2001a), and currently multiple research groups are actively developing marmosets as a model system for studying various sensory and motor systems (Mitchell et al., 2014; Walker et al., 2017), as well as social behavior and disorders (Eliades and Miller, 2017; Miller et al., 2016).

Continued progress of these efforts will depend on neural recording approaches that support the long-term health and well-being of the marmoset while providing chronic access to neural population responses. Current marmoset neurosurgical techniques do not take advantage of the neurosurgical tools available in other species to promote long-term health of the implant and animal (Abe et al., 2009; Adams et al., 2007; Chen et al., 2017; Johnston et al., 2016; Lanz et al., 2013; Parthasarathy, 2014; van Eck and McGough, 2015). Further, restraints typical in work with rhesus macaques, like head immobilization required for tethered recording equipment, when imposed on marmosets have been shown to extinguish voluntary expression of natural behaviors (e.g. vocalization and foraging), limiting the range of possible available avenues for application and investigation (Eliades and Wang, 2003; Walker et al., 2020). While Wang and colleagues (Eliades and Wang, 2008b; Lu et al., 2001b, 2001b; Mohseni et al., 2005; Roy and Wang, 2012) have pioneered an impressive series of neurophysiological recording techniques with marmosets to address these challenges, a solution for longitudinal neural population recordings with minimal disturbance to long-term health and natural behavior has not been developed for marmosets.

Due to the marmoset’s small size, many approaches to recording their neural activity involve removing much of the animal’s scalp to accommodate neural recording hardware and using bone cement to secure this hardware to the skull (Eliades and Wang, 2008b; Lu et al., 2001b; Roy and Wang, 2012). As a consequence, the animal is left with a large open wound for the remainder of its life and limited options for experimental endpoints. While there are recent examples of wireless neural recording solutions without a base of bone cement in larger primates, mainly rhesus macaque, the size of these wireless systems preclude their use with marmosets (Yin et al., 2014). The size of the marmoset’s head limits the battery size, and the battery requires remounting cycles for recharging. Without designing a quick connecting solution, performing recordings throughout the day and night would necessitate significantly disrupting the marmosets’ natural behaviors.

Here we developed a modular wireless headstage configuration to minimize the need for handling marmosets, nearly eliminate the need for restraint, allow for longitudinal studies and completely obviate the need for head fixation during long duration neurophysiological recordings. When combined with the latest neurosurgical techniques (Abe et al., 2009; Adams et al., 2007; Chen et al., 2017; Johnston et al., 2016; Lanz et al., 2013; Parthasarathy, 2014; van Eck and McGough, 2015), for example doing away with the use of methyl methacrylate, we show our methodological innovation facilitates both healing of the neurosurgical site and allows experimental endpoints other than euthanasia. Finally, we demonstrate the flexibility, longitudinal viability and utility of our methods using recordings of neural population responses across the marmosets’ natural behavioral repertoire including sleep.

## RESULTS

A novel platform for chronic wireless neural population recording from freely moving marmosets We have designed and implemented a novel approach to allow for chronic wireless multineuronal recordings in freely behaving marmosets (Figure 1). Our techniques integrate modern neurosurgical tools to maximize long-term health of both implant and animals with a novel neural recording headstage configuration designed to facilitate ease of use (Figure 1A, B) and allow for recording with multiple marmosets in parallel (Figure 1C).

**Figure 1.**
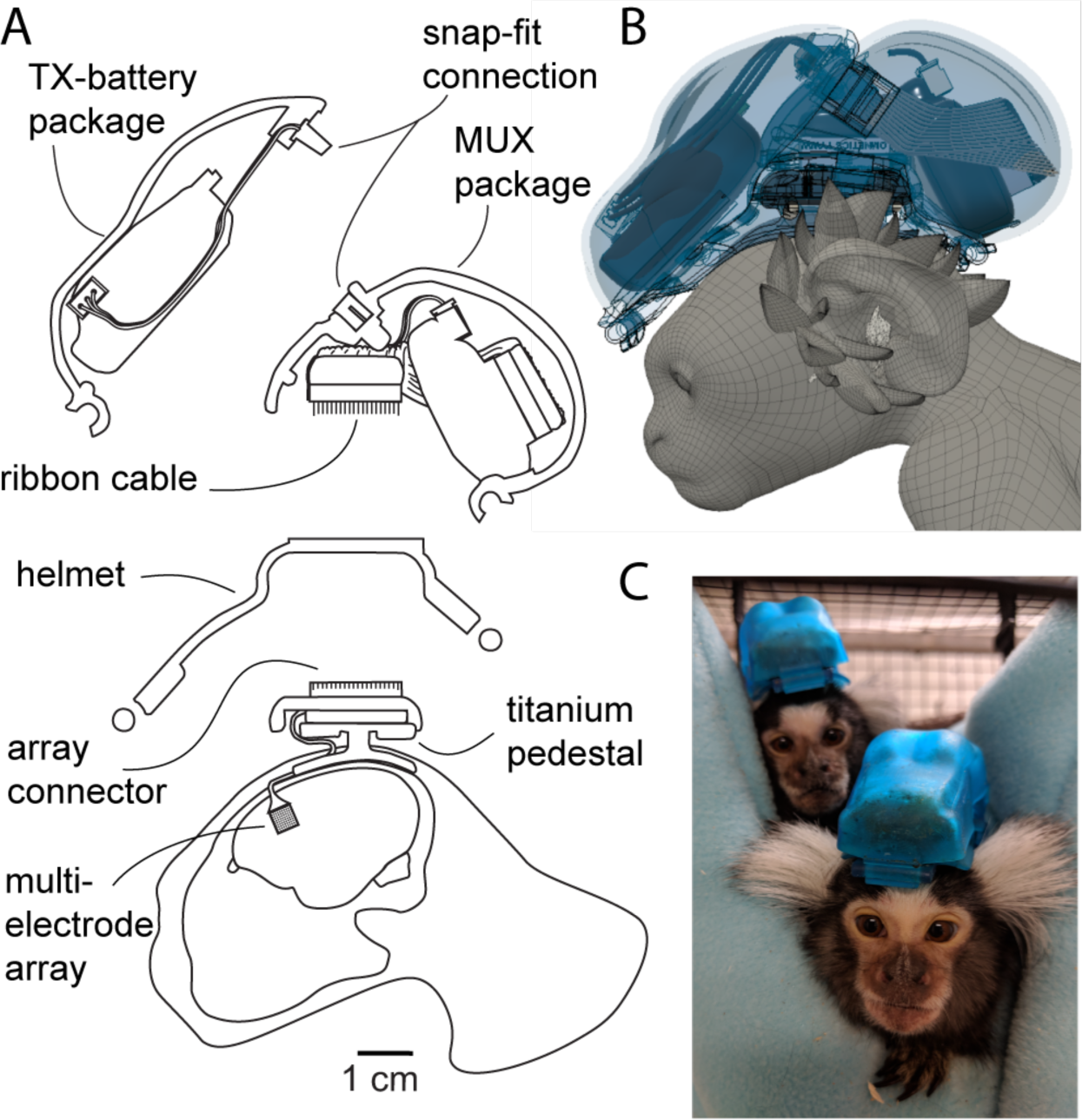
Overview of approach to chronic wireless neural recording with common marmosets. (A) Schematic cutaway illustration of surgical approach components and wireless headstage configuration design. (B) Rendering of headstage assembly. (C) Photograph of two marmosets resting in hammock while wearing headstages for neural recording.

### Surgical approach to implanting the custom titanium pedestal and Utah array

We took advantage of recent advances in human neurosurgery such as custom fit titanium orthopedic implants and ceramic biomaterials to enhance implant integrity and vitality. The resulting surgical preparation keeps the soft tissue of the head relatively undisturbed and accomplishes a 96-channel chronic connection to marmoset sensorimotor cortex with just a 3 mm diameter transcutaneous element and no methyl methacrylate cement. These cements can lead to implant instability and infection in work with macaques, and have fallen out of favor as a result (Adams et al., 2007; Johnston et al., 2016; Lanz et al., 2013). To maximize the potential for long-term animal health and array longevity, we designed a custom, form-fitting titanium pedestal to secure the connector for the electrode array to the skull (Figure 2). In addition, we use multiple preparations of hydroxyapatite, a ceramic material similar in composition to bone known to increase biocompatibility, to further refine the preparation (Figure 2 circular detail insets). As a result, we have been able to record population activity from the sensorimotor cortex of two marmosets for multiple years.

**Figure 2.**
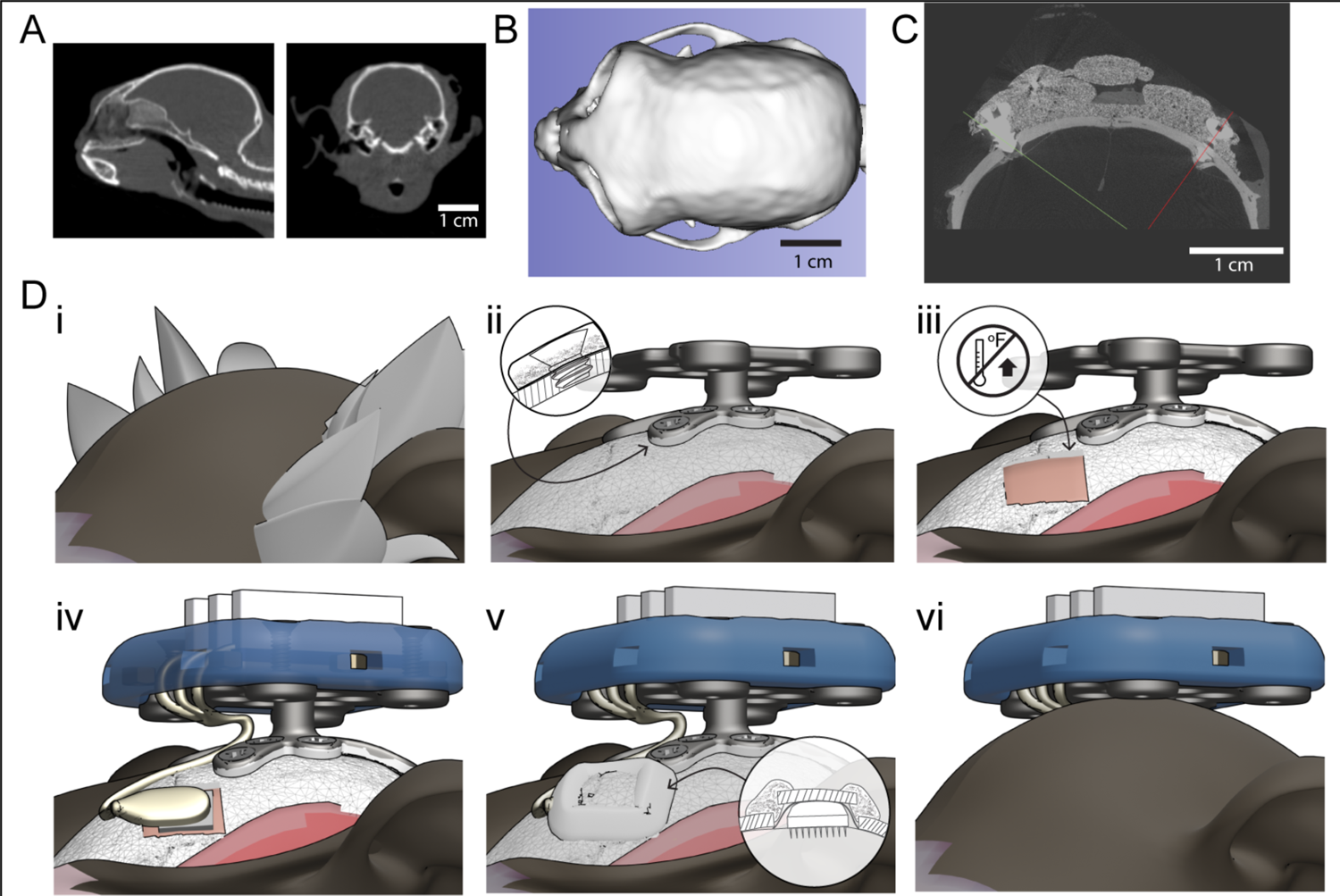
Anatomical considerations and surgical technique for maximizing long-term health for chronic electrode array implantation. (A) CT scan of marmoset cranium, sagittal and coronal sections showing skull thickness and temporalis insertions. (B) Dorsal aspect of 3D marmoset skull segmented from CT scan. (C) Coronal section of marmoset calvarium CT scanned after more standard surgical approach with bone screws to anchor a mound of methyl methacrylate cement used to fix neural recording hardware to the marmoset skull. Note large cranial footprint of central cement mound with two bone screws (indicated with green and red lines) penetrating parietal bone on either side of cement. (D) Surgical stages for implanting Utah array using refined surgical techniques that replace methyl methacrylate cement with custom fitting titanium pedestal for securing neural recording hardware to the skull. Procedure outline: i) pre-surgery, ii) install titanium pedestal with hydroxyapatite coated feet and careful screw placement, iii) prepare craniotomy taking care to minimize thermal accumulation, iv) insert multi-electrode array, v) close craniotomy with artificial dura, native bone and/or hydroxyapatite bone substitute, vi) suture midline incision around wire bundle and pedestal stalk.

Several aspects of marmoset cranial anatomy constrain the design of hardware to secure neurophysiological recording equipment. First, the marmoset occipital, frontal and parietal bones are no more than 1 mm thick (Figure 2A, C). Second, the location of the temporal line relative to the sagittal suture, is variable across marmosets (between 5 – 10 mm lateral to the midline) and constrains the size of the surgical field available before needing to resect the temporalis muscle (Figure 2A, B). If the heads of the cranial screws are not embedded in cement, the installation of cranial screws, either as grounds or for fixation, must be done precisely to avoid penetrating the cranial vault (Figure 2C). The geometry of the pedestal base we designed respects the temporalis insertions and eliminates the need to resect them during surgery. The slender stalk of the pedestal (Figure 2D) creates just a 3 mm transcutaneous object, as opposed to a transcutaneous mound of bone cement many times that size. Further, there is no appreciable sub cranial space in marmosets. As a result, unless the cranium is left open, a solution for cranial closure must be employed after implanting electrodes. We designed a surgical approach that addresses these anatomical considerations and prioritizes implant longevity and animal health.

To implant the electrode array and titanium pedestal for securing the array connector, we made a midline incision and retracted the skin laterally to expose the frontal and parietal bones (Figure 2D i and ii). We then used a custom drill guide to prepare pilot holes for the titanium screws for fixing the pedestal to the skull (Figure 2D ii, Supplementary Figure 2). The geometry of the pedestal platform then served as a guide for the installation of titanium screws taking care to apply even pressure and turn the screw only enough to seat its threads a depth of 1 mm or less (Figure 2D ii inset).

Once the pedestal was installed (Figure 2D iii), we prepared a 6 x 7 mm craniotomy centered on the stereotaxically defined center of the forelimb area of primary motor cortex (Burman et al., 2014, 2008). We used a pneumatic drill with a 0.5 mm rose burr to prepare this craniotomy, taking great care to minimize thermal damage. We placed the resulting bone section in a bowl of cold sterile saline. With the dura intact, we iteratively bent the gold wire bundle until the electrode tips sat on a plane tangent to the cortical surface when the array connector was in place on the pedestal. After the wire bundle was bent appropriately, we removed the dura using a 26 G needle with a tip bent into a hook. We hooked the dura in one corner of the craniotomy site and used either another 26 G needle or fine iris scissors to resect the dura on three sides of the craniotomy to expose the brain.

With the brain exposed, we stimulated the cortical surface to confirm motor cortex location by eliciting movements of the forelimb. After surface stimulation, the array connector was secured to the pedestal with a 3D printed connector fixture (Figure 2D, iv). The Utah array was configured with two reference/ground wires exiting the connector and two integrated reference wires exiting the array board perpendicular to the recording electrodes. The two wires exiting the connector were secured to the pedestal screws. The two exiting the array board were trimmed to fit within the boundaries of the craniotomy. After confirming the electrode tips were still flush with the cortical surface, we inserted the electrodes into the cortex with a pneumatic inserter.

When implanted, the array board sat flush with the skull surface and the wire bundle confluence above it sat above the surface of the skull (Figure 2D iv and v inset). To close the craniotomy, we covered the array with a layer of artificial dura and tucked the edges of this material under the native dura on the margins of the craniotomy as much as possible (Supplementary Figure 3A). We placed the section of bone removed when preparing the craniotomy on top of this artificial dura. Due to the thickness of the wire bundle and silicon insulation leaving the array, and to the kerf of the rose burr used to prepare the craniotomy, there was about a 1 mm gap between the bone section and the edges of the craniotomy (Figure 2D v inset). We either used a hydroxyapatite bone substitute to seal this gap and close the craniotomy (Figure 2D v), or we did not replace the bone flap and used the artificial dura and bone substitute alone to close the craniotomy (Supplementary Figure 3A). Once this bone substitute was set, we sutured the midline incision such that its margin was as close to the pedestal stalk as possible to minimize the size of the cutaneous disruption (Figure 2D vi).

**Figure 3.**
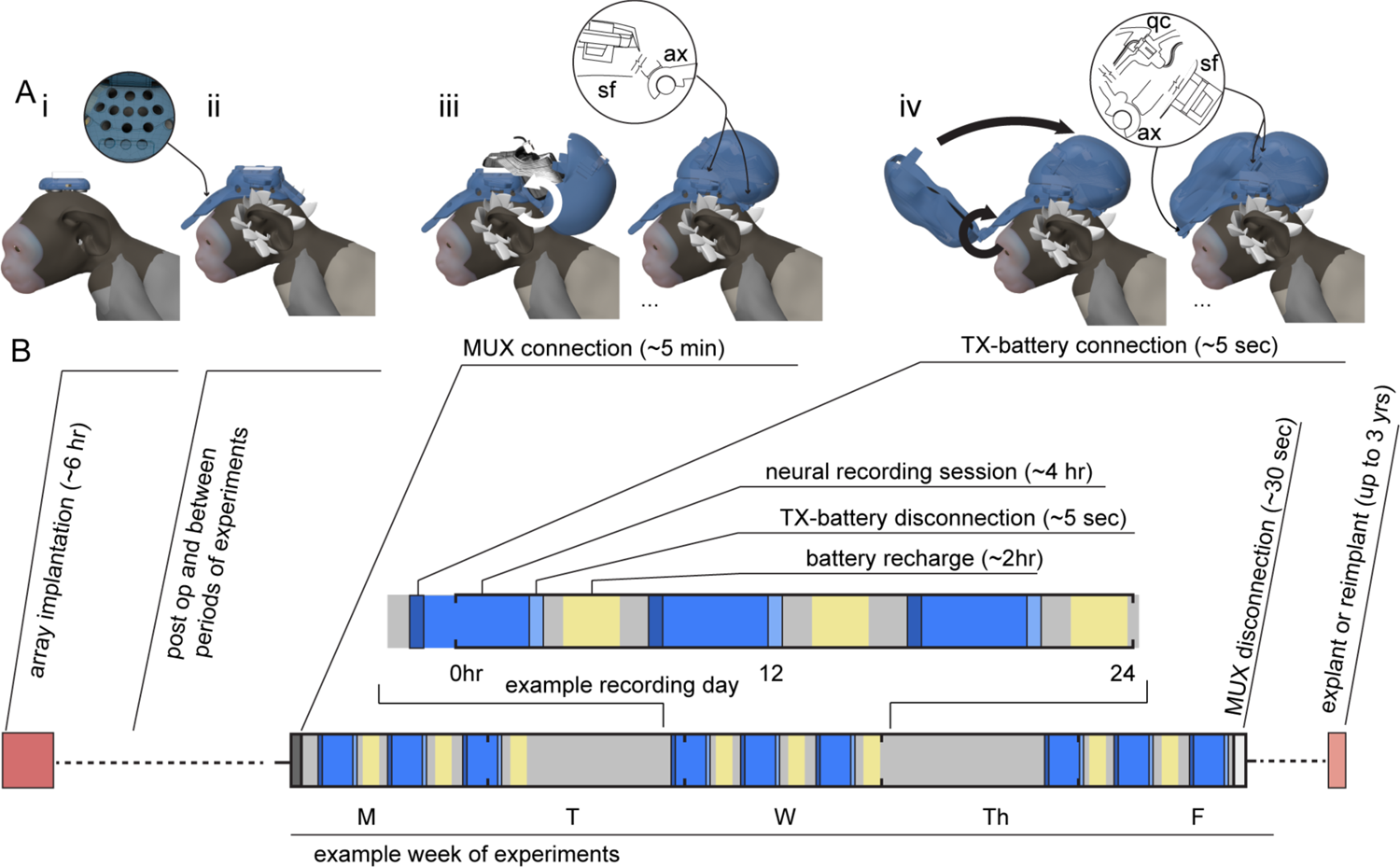
Modular, easy to assemble wireless headstage configuration facilitates recharging battery without disrupting natural behavior. (A) Postoperative assembly with array connector secured to pedestal (i) and helmet (ii) protecting connector and craniotomy site with perforations to allow visual inspection and air flow. (ii) Array assembly when not performing daily recordings. (iii and iv) Head mounted assembly during experiments (e.g. when we are performing neural recordings daily; colored grey in timeline) with MUX package connected to array with 32-channel ribbon cables and 3D printed snap-fit housing mounted to protect the MUX package. (iv) Assembly during neural recordings (colored blue in timeline) with TX-battery package attached to MUX package with three-pin connector and snap-fit housing for TX-battery package. Three-pin connection is made automatically by kinematic constraints designed into the housings. Curved arrows indicate direction of motion for attaching housings. Insets detail elements of kinematic constraints that facilitate expedient assembly: sf, snap-fit; ax, axle; and qc, quick connect. (B) Illustration of experimental timeline detailing sequence of steps for headstage operation during an example period of multiple daily experiments (grey), including neural recordings (blue) and recharging the headstage battery (yellow).

Finally, a 3D printed helmet was attached to the pedestal assembly to protect the craniotomy, the sutures and wire bundle (Figure 3A i to ii). This helmet later served to protect the surgical site and as the platform on which the components of the wireless headstage sat during neural recordings.

### Assembly of easily connectable modular headstage and housing for recording

To facilitate neural recordings, we designed a custom modular configuration of the W-64 wireless headstage (Triangle Biosystems Inc) that permits minimal handling of the marmosets for operation. The marmoset’s small size, finite battery capacities and connectors with many small pins posed the primary challenges to creating a solution for easy to use, chronic wireless neural recordings. We designed a headstage configuration that allowed us to record across the marmoset’s natural behavioral repertoire, and during sleep, with minimal disruption to those behaviors (Figure 3). As battery life is finite and the marmoset’s small size limits battery size, any existing battery needed to be removed, recharged and replaced for longitudinal recordings. Further, the Utah array connector solution intended for large primates is too large for the marmoset head, and the alternative is a PCB with Omnetics nano connectors. To seat the wireless headstage to these connectors, 80 of 120 small, tightly packed pins must be precisely aligned. The precision and force required to make this connection necessitates immobilization of the marmoset’s head.

The headstage configuration we adopted was our solution to avoid repeated immobilization of the marmoset for battery removal and replacement while performing neural recordings across the marmoset’s behavioral repertoire and during sleep. The headstage configuration is split into two packages: a multiplexer and accelerometer package (MUX package, 11 g and 23 x 33 x 15 mm) that was worn on the back of the head, and a transmitter and battery package (TX-battery package, 13 g and 15 x 50 x 15 mm) that was worn on the front of the head (see Figure 1). The MUX package connects to the array connector through two 32-channel Omnetics nano connectors and stays on the head of the animal throughout periods when multiple recordings will be performed (Figure 3A iii and indicated in grey in Figure 3B). The MUX package reduces the number of wires needed for transmitting neural signals to three and conveys them through a three-pin snap-fit connection to the TX-battery package for wireless transmission to the recording PC (Figure 3A iv inset). This three-pin snap-fit connection is made simply through motion constraints imposed by the design of the headstage housing (Figure 3A iv and indicated in blue in Figure 3B). The motion constraints and snap-fit design make it possible to remove and replace the battery for recharging (indicated in yellow in Figure 3B) without picking up the marmoset, using positive reinforcement or while in their hammock. As a result, we could record from multiple marmosets with minimal disturbance to the marmosets’ natural behavior.

To illustrate the advantages of this headstage configuration, we provide a timeline of an example week during which we conducted multiple daily neural recording sessions (Figure 3B). The first mode of headstage operation started at the onset of a period during which we performed repeated neural recordings (3A iii MUX connection, indicated in dark grey in Figure 3B). We attached the MUX package to the back of the helmet and connected its ribbon cables to the array connector. These ribbon cables consisted of Omnetics nano connectors bringing neural signals from the array connector to the multiplexer. Mating these connectors did require briefly holding the marmoset. Critically, this only happened at the beginning of a period during which multiple recording sessions would take place, rather than before and after every recording session. The most posterior element of the helmet was an axle to which a hinge element on the housing for the MUX package attached. The housing rotated forward around this axle to enclose the MUX package and engaged snap-fits on either side of the helmet to secure the MUX package assembly to the helmet.

The second mode of operation began when we attached the TX-battery package and actually performed neural recordings (Figure 3A iv and indicated in blue in Figure 3B). This was done with a similar hinged snap-fit design with the addition that engaging the snap-fits also mated the three-pin connection between the MUX and the TX-battery package (Figure 3A iv inset, qc). The constraints imposed by the hinge on the motion of the housing made this three-pin connection automatically. Consequently, there was no need to handle any wires to make this connection. As a result, to attach the battery, the marmoset’s head only needed to be somewhat stationary for just a few seconds. With adequate positive reinforcement or proper timing (when resting in a hammock), the battery could be attached and removed without handling the marmoset at all.

### Use Case A: Recording sensorimotor population activity and kinematics during natural foraging behavior

Coupling precisely quantified kinematics of the upper limb with recordings of populations of neurons in premotor, primary motor, and primary somatosensory cortices has led to an understanding of cortico-motor function and the development of neuroprosthetic devices to restore lost motor function (Lebedev and Nicolelis, 2017). Using a semi-automated behavioral training approach (Walker et al., 2020), and an approach to quantify kinematics using bi-planar X-ray (XROMM) (Brainerd et al., 2010a), we could train marmosets to engage in ethologically relevant unrestrained upper limb tasks and record kinematics of that behavior. These capabilities, coupled with the approach to wireless neural recording we described above, allowed us to record sensorimotor cortical population activity simultaneously with upper limb kinematics while marmosets engaged in foraging (Figure 4).

**Figure 4.**
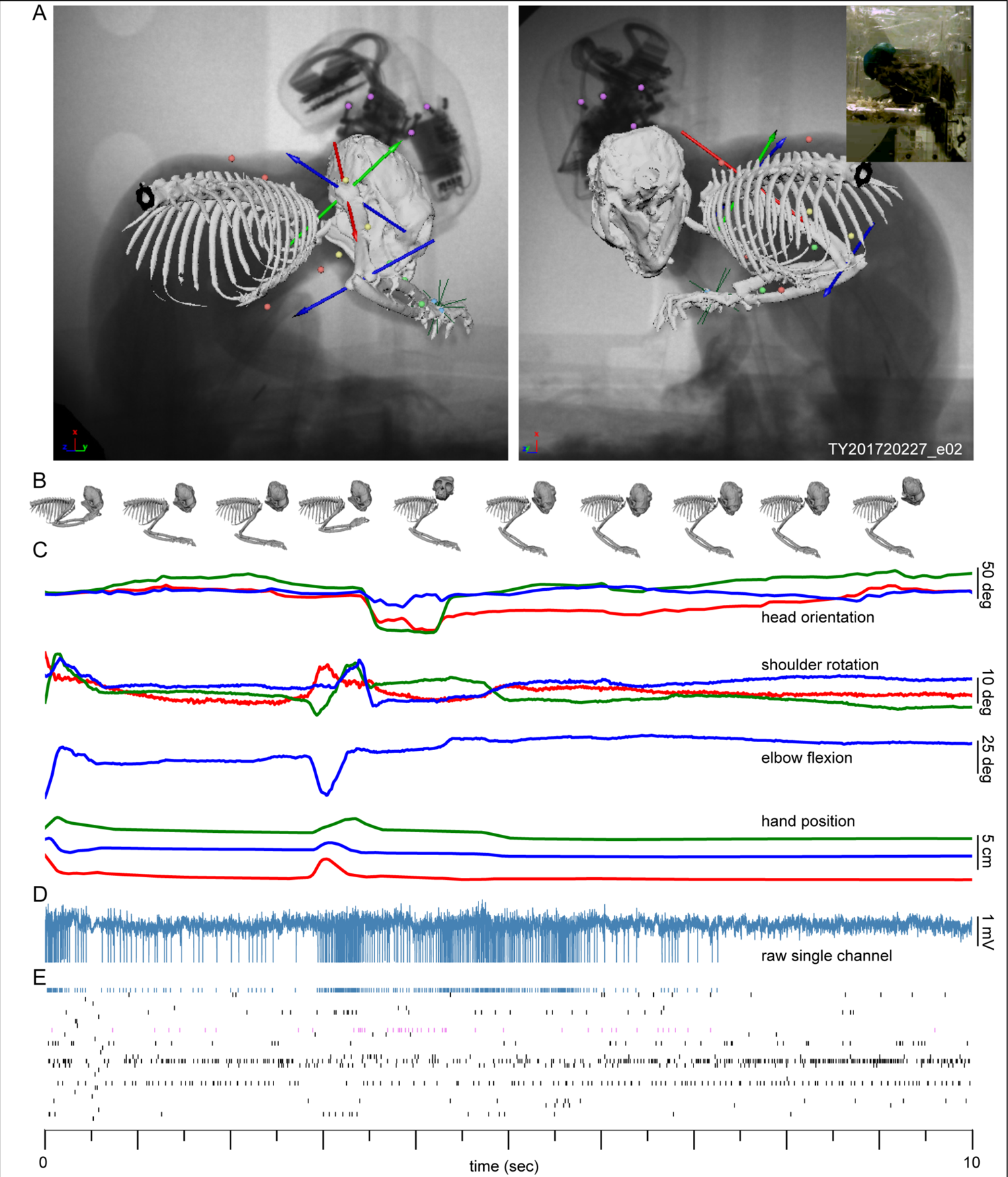
Recording neural population activity simultaneously with upper limb kinematics during foraging. (A) Single frame from of X-ray video capturing foraging behavior with the XROMM. Note colored circles indicating subcutaneous radio-opaque markers placed in the arm (green, forearm; yellow, humerus; blue, hand; and red, torso) which are tracked to reconstruct kinematics. Also note axes

Using bi-planar X-ray, we tracked a combination of small subcutaneous radio-opaque beads (Figure 4A, colored circles) in the torso, upper arm, forearm and hand, and fiducial markers in the pedestal assembly to compute rigid body transformations of the head, torso, humerus, forearm and hand. From these rigid body transformations, we reconstructed the relative motion of these bones (Figure 4B) and then defined joint coordinate systems (Figure 4A, axes) for quantifying shoulder and elbow motion (Figure 4C). The rotation of the rigid body computed with the head markers provided head orientation. As a step toward identifying relationships between upper body kinematics and sensorimotor cortical population activity, we simultaneously recorded neural signals (Figure 4D-E). X-ray generation required for capturing kinematics did not significantly interfere with signal quality of wireless recording of neural signals (Figure 4D). As expected, population activity was heterogeneous; however, relationships between single unit responses and upper limb kinematics and head orientation were readily apparent (Figure 4E, highlighted in blue and violet respectively).

### Use Case B: Recording sensorimotor population activity across the marmosets’ unconstrained behavioral repertoire

The foraging behavior described above takes place within a broader repertoire of natural marmoset behaviors. A consistent finding from a recent body of work investigating motor cortical activity from a dynamical systems perspective is that population activity can be represented in fewer dimensions than the number of recorded neurons (Gao and Ganguli, 2015). This fact has been shown to have functional implications for movement preparation, execution and learning (Afshar et al., 2011; Kaufman et al., 2014; Sadtler et al., 2014).

However, it remains unclear to what extent this finding of reduced dimensional dynamics might be attributed to the use of a limited set of simple tasks. Further, the extent to which insights gained during simple tasks transfer to more unconstrained behavior which require postural control for instance is unknown and will have implications for the development of neuroprosthetic devices that function outside of controlled laboratory settings. As a step toward understanding motor cortical dynamics during unconstrained natural behavior we used our methodology to record sensorimotor cortical population activity during unconstrained natural behavior.

To sample sensorimotor cortical population activity across the marmoset’s natural behavioral repertoire (Figure 5), the wireless transmitter was mounted and the marmoset engaged in a diversity of behaviors within its home cage (Figure 5A-C). Gross behavior was manually annotated using Elan (Brugman and Russel, 2004) and consisted of 14 behaviors (Figure 5D), with six of these behaviors accounting for more than 90% of this hour-long recording: leaping, locomotion, foraging, food manipulation, vertical clinging, and sitting.

**Figure 5.**
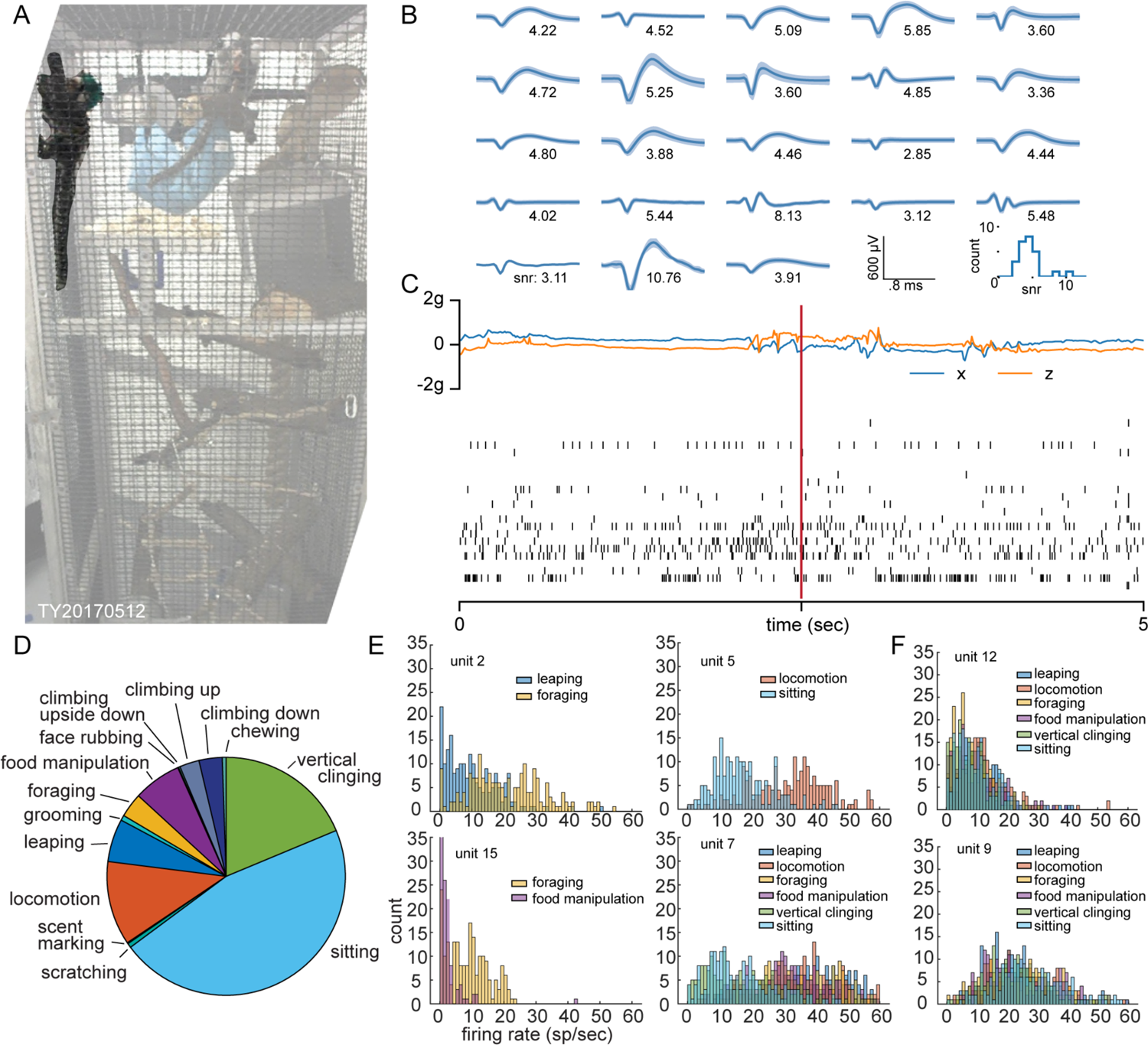
Wireless recording of sensorimotor cortical population activity across the marmoset natural behavioral repertoire. (A) Single frame of video during wireless neural recording of unconstrained marmoset engaging in natural behavior. Note marmoset with headstage assembly vertically clinging to the wall of the home cage in the top left of the image. (B) Mean spike waveforms of neurons from this recording. (C) Accelerometer traces and raster plot of population activity from a short segment of this recording. Vertical red line indicates time of video frame. (D) Summary of one hour of unconstrained behavior annotated with gross behavioral states. (E) Firing rate distributions of example units that exhibited state specific activity. (F) Firing rate distributions of example units that did not exhibit state specific activity.

Evaluating the units in relation to these behavioral states, showed some state-specific rates for two different behavioral states like leaping vs foraging, locomotion vs sitting, and foraging vs food manipulation as well as state dependency on all six behaviors (Figure 5E). We also found units which did not exhibit state-specific firing rates at all (Figure 5F).

### Use Case C: Recording neural activity during sleep

Memory consolidation during sleep has been shown to play a role in motor skill acquisition and learning (Diekelmann and Born, 2010; Gulati et al., 2017; Yang et al., 2014). However, population level understanding of the underlying processes in primates has been limited in part because of the difficulty inherent in facilitating relaxed prolonged sleep during head fixation. Preliminary investigations of marmoset sleep (Crofts et al., 2001; Hoffmann et al., 2012) found marmosets exhibit NREM and REM sleep stages familiar from other primates which cycle with a period of 20 - 40 minutes. To our knowledge, only a single study, recording from auditory cortex, investigated spiking activity during marmoset sleep (Issa and Wang, 2013). With our quick connect configuration of the modular headstage and proper timing we could remove and replace the battery without handling the marmosets. Using this approach, we recorded both single units and LFPs from sensorimotor cortex during sleep (Figure 6A, B).

**Figure 6.**
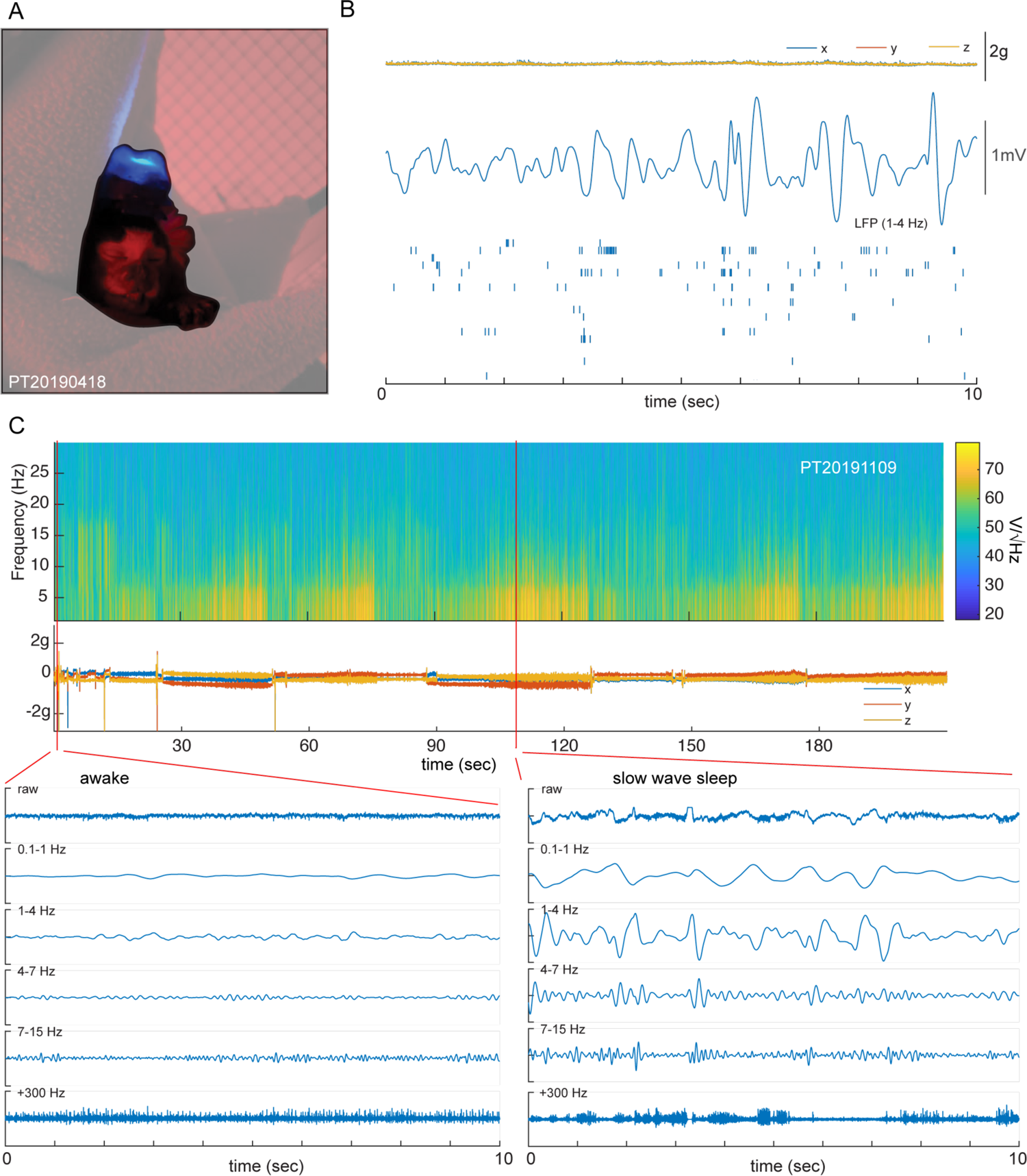
Wireless recording of sensorimotor cortical population activity during sleep. (A) Photograph of marmoset sleeping in hammock while recording cortical population activity. Blue light is the indicator LED of the transmitter. (B) Accelerometer traces, delta band local field potential from single electrode and raster plot from a 10 second segment of this recording. (C) Spectrogram and accelerometer traces from a 3.5 hour long recording of marmoset sleep, with expanded 10 second samples during quite wakefulness and slow wave activity to illustrate cross state differences across frequencies.

We found LFP power across all frequencies below 30 Hz to be low when the marmoset was awake. As the marmoset entered sleep, modulation at low frequencies developed and cycled with a period of 20 - 40 minutes over the course of the recording (Figure 6C). Over the course of a given cycle, we observed increasing low frequency power indicative of slow wave sleep (likely advancing through NREM sleep stages), followed by an attenuation of low frequency LFP power (likely a combination of REM and brief wakefulness). Onset of this transition from high low-frequency LFP power to significantly attenuated low power was often coupled with movement indicated by the accelerometer. Power was distributed through other frequency bands but it was not as strong or as clearly periodic. Finally, we also observed putative sleep spindles in between 7-15 Hz (Figure 6C).

### Array yield over implant duration

With the approaches described above, we were able to record population responses from the sensorimotor cortex of two marmosets for up to three years without any problems with the cranial implants. As expected, there was a gradual decline in the number of single units that could be isolated in the recordings from these arrays (Figure 7). In the case of marmoset PT, we were able to isolate more than 80 units two weeks after implantation, which declined to about 60 units within the first few months after implantation (Figure 7A, B). In the case of the first array of marmoset TY, it began with about 40 units just after implantation and gradually declined to just a few isolatable single units more than two years later (Figure 7B, C). However, perhaps less intuitively, there were instances of units appearing on the arrays where previously there had been none (ex. electrode 9 and 53 in Figure 7A). Additionally, our approach allowed us to replace the failing array in marmoset TY with a new one (Figure 7B blue data point on far right and Figure 8).

**Figure 7.**
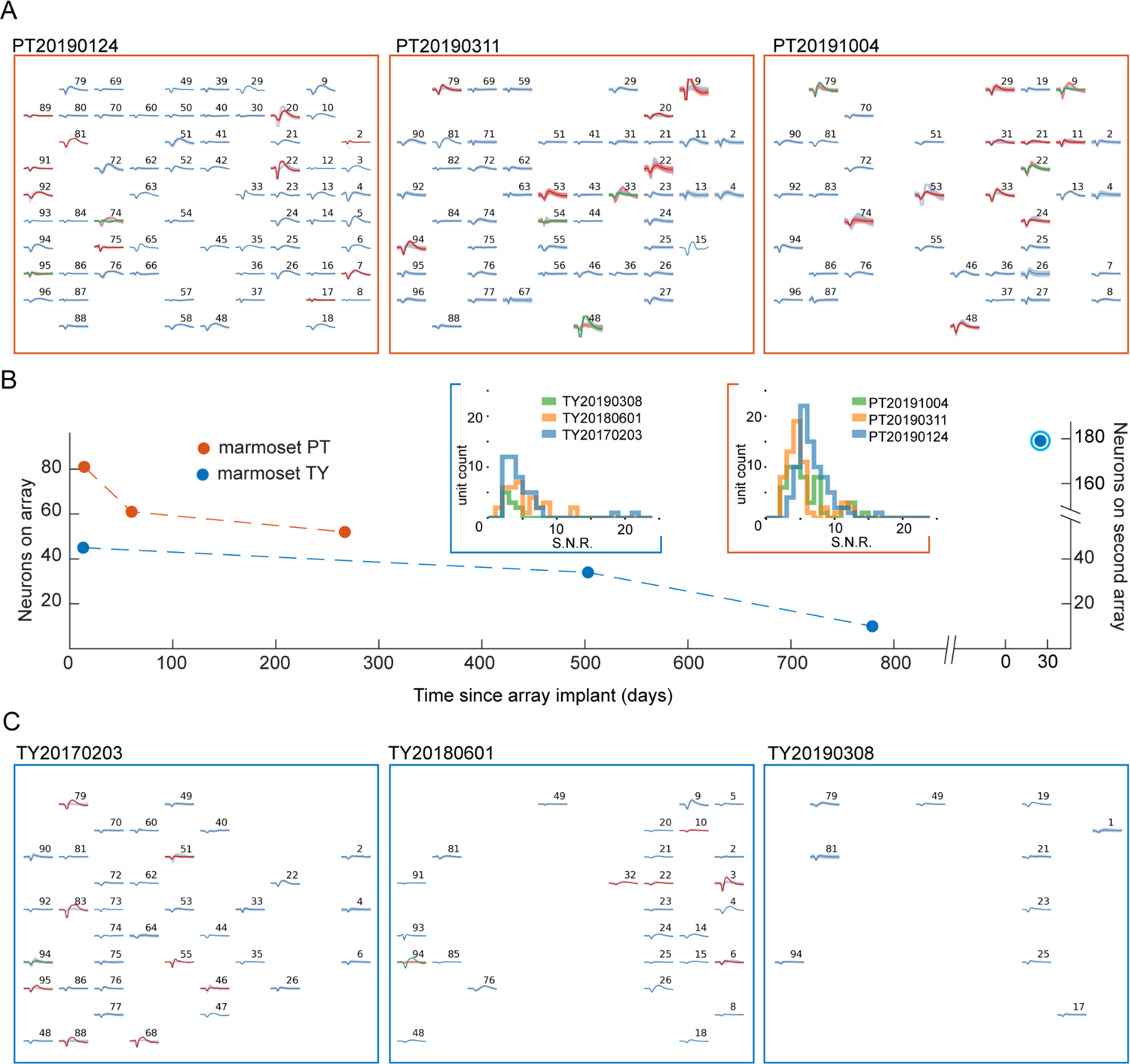
Neuron yield of three electrode arrays after chronic implantation. A) Spike panels displaying mean waveform from three brief recordings distributed over the course of ten months following Utah array implantation in marmoset PT. B) Plot illustrating count of neurons found on each array in the recordings displayed in A and C, with insets illustrating distributions of signal to noise ratios for each of the datasets for both marmosets. Note blue, circled data point on far right illustrating yield of a second array implanted in marmoset TY, also displayed in Figure 8. C) Spike panels displaying mean waveform from three brief recordings distributed over the course of three years following first Utah array implantation in marmoset TY.

**Figure 8.**
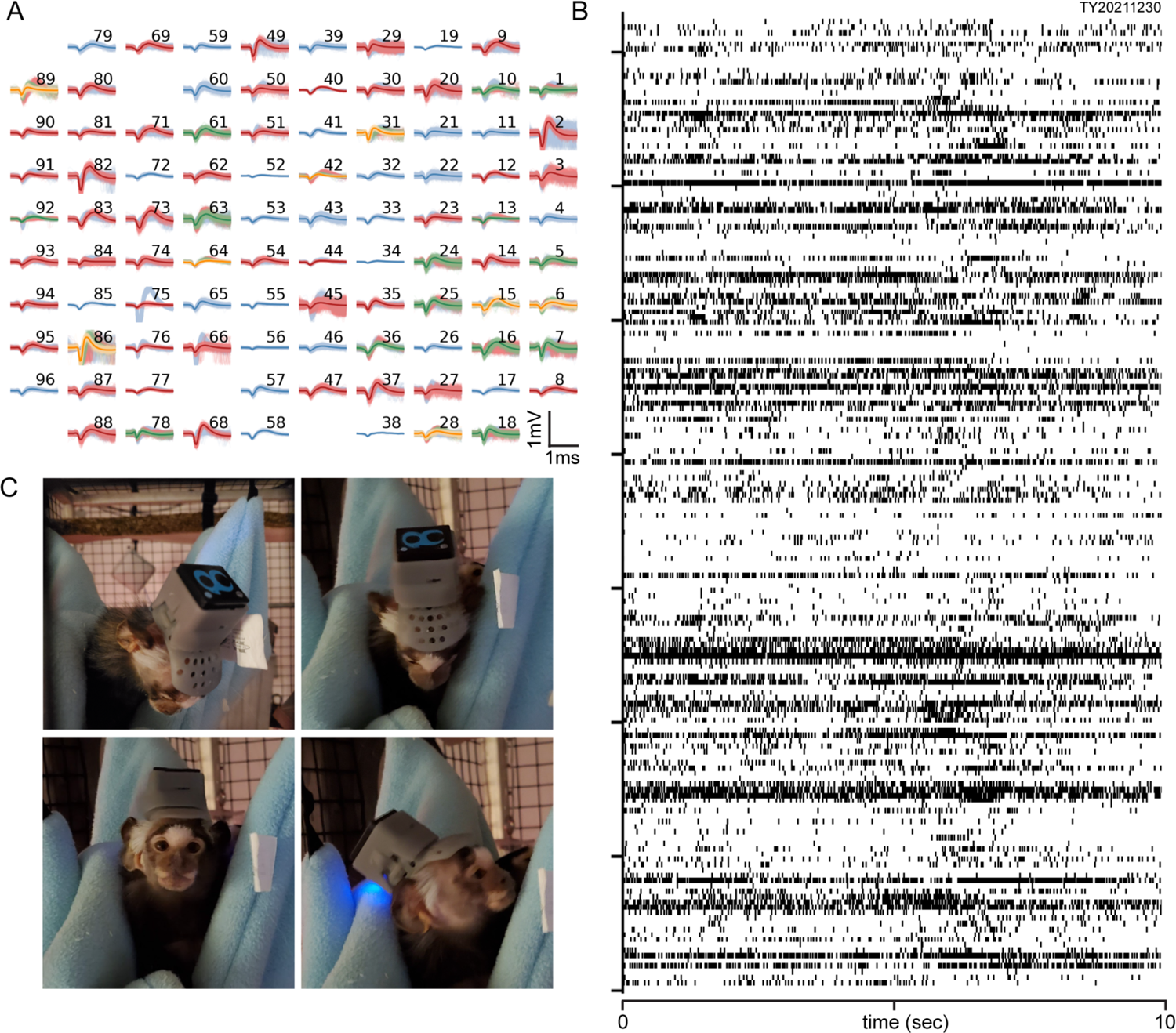
Example data from marmoset’s second array using quick connect approach adapted to a 96-channel wireless headstage. A) Spike panel illustrating mean waveforms and spikes of neurons recorded from a second Utah array implanted in the sensorimotor cortex of the right hemisphere of marmoset TY after removing the connector of the first array implanted in the left hemisphere. B) Raster representing 10 seconds of population activity during free behavior. C) Marmoset TY wearing helmet and 96-channel wireless headstage with quick connect housing.

### Alternative endpoints: Healing and Explant or Reimplantation

Standard approaches to cranial implants with marmosets utilize a mound of bone cement to fix neural recording and head fixation instrumentation (though see Sedaghat-Nejad et al., (2019) for a recent alternative). This often demands a large cranial footprint and necessitates removing substantial portions of the soft tissue on the animal’s scalp. Additionally, these methods often do not replace the cranial bone removed to access the brain. These details mean that the only experimental endpoint available for the animal is euthanasia. In contrast, the custom titanium pedestal with minimal transcutaneous diameter, and sealing of the craniotomy with the autologous bone flap and/or hydroxyapatite bone substitute, allowed for significant healing of the craniotomy (Supplemental figure 5A). Further, as none of the soft tissue of the scalp was removed in surgery our approach offers de-instrumentation as an alternative experimental endpoint. An array that no longer yields usable neural signals can be replaced with a new array using the same titanium pedestal, which we have recently done with great success (Figure 7 and Figure 8) or the titanium pedestal could be removed entirely (Supplemental figure 5B).

## DISCUSSION

Here we designed, refined and implemented an end-to-end solution for performing chronic wireless neural recording from populations of individual neurons in sensorimotor neocortex in unrestrained marmosets. We demonstrated that our approach allows for recording across the marmoset’s behavioral repertoire and during sleep. Our approach includes refined surgical techniques for implanting multielectrode arrays designed to maximize the potential for longitudinal implant, and animal, health while providing the opportunity for alternative experimental endpoints. A key component of our methodology is a modular configuration of the wireless headstage designed to maximize ease of use and minimize disturbance of natural behavior. We have demonstrated that in combination these two methodological advancements allow for longitudinal studies of neural circuit function across the full range of behaviors in which marmosets engage.

Minimizing marmoset handling is a key design feature of the approach. We feel this minimizes marmoset stress, and, coupled with the voluntary behavioral training approach we have developed (Walker et al., 2020), we can perform studies of motor control and learning with minimal disturbance to marmosets’ natural behavior. While the modular headstage assembly we describe was designed around a 64-channel wireless headstage (W-64, Triangle Biosystems), the principles underlying our design, modularity of battery and kinematic constraints for connector couplings toward ease of use, can be applied to other wireless neural recording systems. In fact, we recently adapted the approach we describe here to the 96-channel Exilis wireless headstage (Blackrock Microsystems) to record from a new array implanted in the same marmoset (Figure 7 and Figure 8). In the initial weeks of recording from the new array, we’ve been able to record 1-3 times daily while only lifting the marmoset out of its home cage a few times. Using the behavioral training approach we have described previously (Walker et al., 2020), this marmoset has also been voluntarily engaging in experimental reaching tasks and completing hundreds of reaches per session. The combination of these two approaches amounts to a very productive experimental setup for studying the neural coordination of voluntary movement in marmosets without any food or water restriction and no daily restraint.

We have developed an end-to-end solution for performing chronic wireless neural population recordings with marmosets in behavioral neuroscience. Our refinements of the electrode implantation procedure, the use of a custom, form-fitting titanium orthopedic implant and multiple preparations of hydroxyapatite bone substitute, offer opportunities for osseointegration, improved healing, electrode replacement and alternative experimental endpoints. These design features enable chronic recordings and longitudinal studies of skill acquisition. Coupled with the quick-connect wireless headstage configuration, the techniques we present open new avenues for the study of neural circuit function during unconstrained behaviors across the marmoset repertoire of natural behavior, including social behavior and sleep.

### EXPERIMENTAL MODEL AND SUBJECT DETAILS

#### Common marmosets, *Callithrix jacchus*

The results and methods described in this paper were obtained with two adult Common marmosets (one male and one female, ages 7-8, 350 - 415 g). Neither marmoset had been involved in any previous studies and both were healthy during the duration of the work reported in this paper. However, one of the marmosets did ultimately succumb to marmoset wasting syndrome, an inflammatory bowel disease, and had to be euthanized. These two marmosets were pair housed in a 2’ x 2’ x 6’ home cage furnished with branches, a nestbox, a foraging shelf and a hammock. Diet included ZuPreem Marmoset Diet supplemented daily with a variety of dried and fresh fruits and nuts. The light cycle in the housing room was 12 hours on, 12 hours off, and the temperature was set between 78 and 86°F with a target humidity of 30 – 70%. All surgical procedures and animal handling methods were performed in an AAALAC accredited facility in accordance with the standards described in the Guide for the Care and Use of Laboratory Animals (National Research Council, 2011), and were approved by the Institutional Animal Care and Use Committee of the University of Chicago.

### METHOD DETAILS

#### Design and fabrication of the custom titanium pedestal

Three main factors constrained the design of the pedestal geometry: marmoset cranial anatomy, the size of the small animal connector for the Utah array (Blackrock Microsystems), and the requirement to maximize longitudinal implant health. The design process began with a micro-CT scan of a marmoset skull (150 kV, 150 µA, 33 ms exposure) using the 240 kV directional focus tube in the paleoCT facility at the University of Chicago. This scan resulted in a Dicom file (.dcm), which was segmented using Amira (ThermoFisher Scientific) to extract the geometry of the marmoset skull as an .stl file. This file representing the marmoset cranium was then imported into Fusion 360 (Autodesk) for all subsequent design and surgical planning. In addition to the marmoset skull, we also manually created a 3D model of the small animal Utah array in Fusion 360. We created a stereotaxic coordinate system, to match that described by Paxinos et al., (2012) and located the boundaries of the craniotomy that would be performed in surgery.

To model the pedestal, we created a sketch plane tangent to the cranial surface above the sagittal suture and sketched the outline of what would become the feet of the pedestal being careful to respect the temporal lines on the parietal bones. We then created a t-spline plane and used the point snapping function to make the curvature of this plane match the curvature of the cranial surface. We extruded the sketch plane representing the pedestal feet toward the t-spline plane matching the curvature of the skull and kept the intersection, which was then the geometry of the feet with curvature matching that of the skull. We then thickened this to 1 mm and added a 3 mm diameter, 3.6 mm high cylinder up from the center of the feet. We softened all the sharp edges with 0.5 - 0.9 mm fillets and added a 0.6 mm equal distanced chamfer to the screw holes to match the 90° angle of the heads of the screws that were used to fix the pedestal to the skull. We then added a 1 mm thick platform large enough to accommodate the connector for the electrode array and one screw on each corner for fixing the connector to the pedestal. We thickened the portions of the platform to later accommodate 0-80 threading, removed material of this platform to accommodate hand tools for installing the cranial screws in surgery, and rounded sharp edges.

The geometry of the pedestal was exported from Fusion 360 as an .stl. This file was sent out for fabrication in titanium (Ti64, proto3000, Ontario, Canada) using direct metal laser sintering (DMLS). Upon receipt of the fabricated titanium pedestals, we manually filed the convex surface of the feet using a needle file to remove any roughness left by the supports used for DMLS fabrication. We also refined screw holes with a drill press to clean up any roughness, before manually threading the through holes on the platform with a 0-80 tap. After the pedestal was cleaned of any roughness and its holes were tapped, it was sent to have its feet coated with hydroxyapatite through plasma spray (APS materials, Inc., Dayton, Ohio) to promote biocompatibility and osseointegration.

We designed a fixture to hold the array connector PCB onto the platform of the pedestal. The fixture began as a sketch on the top surface of the pedestal platform that framed both the platform and the array connector. A series of extrusions was made to create a 5 mm thick form that sandwiched the PCB on the pedestal platform and only exposed the three Omnetics nano connectors used to gain access to the electrode signals. The connector was fixed to the pedestal with 0-80 7/32” Philips head machine screws installed in the threaded holes of the pedestal platform. In addition, this fixture held six captive brass nuts (part no. 92736A001, McMaster Carr) which were used to attach the protective helmet to the fixture after the array implantation.

#### Surgical procedure for implanting the pedestal and electrode array

Prior to surgery, all surgical tools and components for implantation were sterilized by autoclave or ethylene oxide. Marmosets received a single intramuscular (IM) dose of dexamethasone (0.5 - 2 mg/kg) the afternoon before and were fasted 12 hours prior to surgery, during this time apple juice was provided to prevent potential hypoglycemia. The morning of the surgery, marmosets were given an additional dose of dexamethasone (0.5 mg/kg), which was tapered over the next few days, and antiemetic maropitant citrate (1 mg/kg). A single IM dose of ketamine (10 mg/kg), or Alphaxalone (5 – 10 mg/kg), was given for anesthesia induction, and anticholinergic atropine sulfate (0.02 – 0.04 mg/kg) was given to reduce excessive salivation and prevent bradycardia during intubation. Marmosets were then intubated with an uncuffed murphy eye endotracheal tube (ID 2.0 or 2.5 mm) and given isoflurane, 1 - 3 %, to maintain a surgical plane of anesthesia. Their head, leg and arm were shaved and scrubbed with three cycles of betadine, with rinses of 70% isopropyl alcohol, and an intravenous catheter (24G) was inserted into either the saphenous vein or the lateral tail vein. Their head was secured within a head holding adaptor (CA-6, Narishige) for the stereotaxic frame. Marmosets were then placed in a warmed stereotaxic frame (SR-6C-HT, Narishige). Throughout the procedure the following vital signs were monitored continuously: heart rate, EKG, end tidal CO2, blood oxygen saturation, rectal temperature, blood pressure and respiratory rate. Additionally, if CO2, oxygen saturation and respiratory rates fell outside of normal ranges, the marmosets were ventilated with positive pressure using tidal volumes of 10 – 20 mg/kg and a rate of 8 – 20 breaths/minute, with a pressure-controlled ventilator (Engler ADS 2000) to prevent any damage to the lungs from overventilation.

A midline cranial incision was made from slightly posterior to the glabella to the lambdoid suture. Skin was parted and held laterally by sutures tied off at the four corners weighted by light hemostats. A stereotaxic manipulator arm, zeroed prior to placing the marmoset in the stereotaxic frame, was mounted to the rail of the stereotaxic frame, and adjusted to mark the location of stereotaxically defined forelimb M1 (9.8 AP and 4.5 ML). The intersection of the DV line at this point with the cranial surface marked the center of the craniotomy. We then measured out from this line 3 - 4 mm to mark the boundaries of the craniotomy with a sterile pen.

With the boundaries of the craniotomy indicated, the titanium pedestal was placed on the cranium such that their respective curvatures matched. With the pedestal held in place, the surgeon drilled a single pilot hole less than or equal to 1 mm in depth with a 1.2 mm drill bit (part # 30565A215, McMaster Carr) held by a pin vise (part # 8455A31, McMaster Carr). A single M1.4 x 2 mm titanium screw was manually installed into this pilot hole through one of the holes in the feet of the pedestal using firm, even pressure and turns of the screw driver (part # 26105, Wiha tools) to match the number of threads that fit within the 1 mm thickness of the parietal bone. The goal was to have the titanium screw threads tap the pilot hole without stripping it.

The pitch of the screw’s threads was chosen to roughly match the thickness of cortical bone layers, as determined by μCT scan, and to have the threads cut between these layers. With the pedestal fixed to the skull using one or two of these pilot screws, the surgical guide was placed onto the pedestal to facilitate maintaining an entry angle perpendicular to the cranial surface and limiting the depth of pilot holes to under 1 mm. The remaining pilot holes were drilled manually using the pin vice and surgical guide.

Once the pedestal was firmly affixed to the skull with cranial screws, we began preparing the craniotomy using a 0.5 mm rose burr (part # 19007-05, Fine Science Tools) and a pneumatic drill (part # 2910-100, Micro-Aire). We minimized thermal accumulation by frequently quenching the rose burr in cold sterile saline, irrigating the craniotomy borders, minimizing repeated passes over the same edge of the craniotomy, and focusing on economy of movement. Probing the boundaries of the craniotomy with a small spatula by applying gentle pressure around the edges of the bone flap and looking for motion of the bone, or lack of motion, provided a good indication of the depth and evenness of the craniotomy preparation.

When motion could be seen around all of each edge of the craniotomy when gentle pressure was applied, the tip of the spatula was placed within the kerf of the craniotomy and used to very gently lift the central bone segment away from the underlying dura. This bone segment was then submerged in cold, sterile saline for safe keeping until after the array was inserted to facilitate closing the craniotomy.

After the craniotomy was complete, but before performing the durotomy, we prepared the electrode array for implantation. First, we trimmed the references and ground wires such that the two exiting the electrode board sat within the boundaries of the craniotomy and the two exiting the rear of the connector PCB were long enough to wrap around the two posterior screws securing the fixture to the pedestal. Second, we trimmed the insulation off the tips of these wires with fine forceps. Lastly, we iteratively bent the gold wire bundle such that the electrode tips sat on a plane tangent to the newly exposed dural surface when the array connecter sat on the pedestal. This pre-positioning of the electrode array before insertion was critical. We choose to do it before removing the dura to protect the brain surface from repeated electrode contact.

With the wire bundle preshaped and the electrode array ready for insertion, we performed a durotomy using a 26G needle with its tip manually bent into a hook and either iris scissors or another 26G needle. The bent needle was used to grab the dura and the iris scissors or unbent needle were used to incise the dura on the anterior, medial and posterior edges of the craniotomy. A section of gelfoam saturated with cold sterile saline was placed on the lateral edge of the craniotomy and the dura was folded up onto it. A second section of saturated gelfoam was placed onto the dura in an effort to keep it moist until closing the craniotomy, though we found that, by the end of the procedure, the dura had shrunk and was not large enough to suture over the array.

Once the brain was exposed, we used a monopolar stimulating probe (RLMSP051, Rhythmlink International) to deliver current to the cortical surface to elicit stimulation effects consistent with primary motor cortex localization (ex. muscle contractions in the upper limb contralateral to the stimulation site). Surface stimulation consisted of 50 Hz biphasic pulses, with a pulse duration of 100 μs, and with currents ranging from 2 – 15 mA (model 2100, AM Systems). The exposed cortical surface was probed by gently touching points with the stimulation probe. While probing the cortical surface, an assistant watched the marmoset’s upper limb for movements or muscle contractions. During this procedure, care was taken to hydrate the cortical surface. Initial stimulations were attempted without modification of the anesthetic regime described earlier. If no stimulation effects were observed, we introduced ketamine in a bolus (1 mg/kg IV), followed by ketamine (15 μg/kg/min) at continuous rate infusion with remifentanil as needed (0.1 - 0.1 μg/kg/min), then, with careful monitoring of heart rate and respiratory rate by veterinary staff, the isoflurane concentration was decreased in 0.25% increments. Surface stimulation had the advantage of being able to cover large sections of the neocortex quickly relative to sharp penetrating electrodes. We also performed intracortical microstimulation as well as tactile and proprioceptive stimulation after the array was implanted to confirm electrode location (Supplementary Figure 4).

After cortical stimulation, we were ready to fix the array connector to the pedestal and insert the electrode array into sensorimotor cortex. A stereotaxic manipulator arm was attached to the stereotaxic frame to hold the actuating wand of the pneumatic inserter (Blackrock Microsystems) using a custom machined adaptor and a ball joint (part # BC-4L, Narishige). The connector for the electrode array was placed onto the platform of the titanium pedestal to ensure that the electrode tips were still flush with the brain surface with the array centered on cortical surface from which stimulation effects were elicited. If necessary, additional bending of the wire bundle could be done to refine the final position of the array before insertion. Once the array sat flush with the cortical surface, and its reference wires were within the boundaries of the craniotomy, the 3D printed fixture (Tough Resin, Formlabs) was added to secure the array connector to the pedestal platform. While installing the screws (0-80, 7/32, part # 91771A040, McMaster Carr) to secure the fixture to the pedestal platform, the reference wires from the connector were wrapped around the screws. With the connector secured to the pedestal and the array positioned on the brain surface, the wand for the pneumatic inserter was placed above the array perpendicular to the cortical surface such that there was no appreciable space between the actuator of the wand and the wire bundle above the array. Once everything was in place, the inserter pump was turned on and the trigger was pressed to actuate the wand and insert the array.

We began to close the craniotomy by attempting to replace the dural flap folded laterally earlier in the procedure. In our experience, this dura will not cover the entirety of the craniotomy and array. To completely cover both the array and craniotomy, we used a section of artificial pericardium (Preclude ePTFE membrane, Gore and Associates, Inc.) cut to the shape of the craniotomy, gently folded over the array and tucked as much as possible under the margins of the craniotomy. The original section of bone removed while preparing the craniotomy was retrieved from the cold sterile saline and placed gently above the layer of artificial pericardium. The marmoset parietal bone is only ∼ 1 mm thick, and once inserted, the dorsal surface of the electrode array, sat ∼ 2 mm above the brain. Additionally, there was a ∼ 0.5 mm kerf left from the rose burr. As a result, there was at least a 1 mm gap between the edges of the bone section and the edges of the craniotomy. To seal this gap and promote the reintegration and healing of the bone section, we sealed this gap with a hydroxyapatite bone substitute (DirectInject, Stryker). This material is similar in composition to bone and is designed to provide a bone growth facilitating medium. We then waited 10 – 15 minutes to ensure this material was adequately set, while preparing to close the midline incision.

Being careful not to disturb the wire bundle, we used monofilament non-resorbable sutures and a simple interrupted stitch to close the midline incision tightly around the wire bundle and the stalk of the pedestal. Once the midline incision was completely sutured, a 3D printed custom helmet (Tough resin, Formlabs) was secured onto the connector fixture with four small screws (part # 91251A054, McMaster Carr) that screwed into the captive brass nuts embedded in the fixture. This helmet served to protect the craniotomy site and the wire bundle. Additionally, this helmet provided a platform for mounting wireless headstage components during neural recordings. With the helmet in place, the marmoset was given buprenorphine (5 – 10 μg/kg) for analgesia and was warmed with a bear hugger (99 - 101° F). The marmoset was monitored to ensure that it could breathe spontaneously. Additionally, adequate blood oxygen saturation (95%) and respiratory rate (40 - 60 rpm) were also verified. Once it was clear that the marmoset was breathing well on its own, the marmoset was extubated and placed on warm soft bedding in a transfer cage. Once the animal was able locomote steadily on its own the IV was removed and it was returned to its home cage with its partner. For 48 – 72 hours after surgery, the marmoset received buprenorphine (5-10 μg/kg) or meloxicam (0.1-0.3 mg/kg) every 12 hours, and antibiotics were given under direction of the veterinary staff. The marmoset was checked multiple times daily and monitored closely over closed-circuit video to ensure there were no signs of pain or distress. Sutures were removed after 14 days under anesthesia.

#### Design of the custom headstage configuration

The default configuration of the W-64 headstage sold by Triangle Biosystems (TBSI) was a single package and provides no solution for mounting or protecting the device on the head of the animal. We worked with TBSI to design a custom modular configuration of the W-64 wireless headstage, where the connection to the array is isolated from that of the battery. As a result, we could leave the connection between the multiplexer and array intact when changing the battery. To take advantage of this feature, we designed a set of nested snap-fit housings to facilitate the connection between the multiplexer and battery packages and protect the headstage assembly. This arrangement greatly minimized the need to handle the marmoset when removing and replacing the headstage battery for charging.

We designed the headstage housing using Fusion 360 (Autodesk) to be 3D printed in Tough Resin (Formlabs) on our Form 2 (3D printer). It began with adding to the helmet additional axle and snap-fit elements designed to facilitate and secure the headstage assembly. The axle elements were added to the rostral and posterior edges, and beams for snap-fits were added lateral to the array connector over the ears. The silhouette of the two headstage packages and their connectors served as a guide for the initial extrusions of their respective housings. To these basic forms we joined complementary clasp and snap-fit elements to constrain the motion of the housings to rotate and snap into place securely onto the helmet. With the kinematics of assembly adequately constrained, we modeled mounting elements within each housing to hold the components of the MUX to battery connection such that this connection was made when securing the snap-fits.

#### Electrophysiological data acquisition

For each recording with the wireless headstage, two of three Omnetics connectors from the array connecter were chosen. To these, the Omnetics ribbon cables from the MUX package were attached. Neural signals from the array were multiplexed with signals from the accelerometer mounted to the rear of the head. These multiplexed signals were then conveyed to the transmitter on the front of the head through three-wire Molex picoblade cable assemblies (parts: 151340400 and 530470310). These signals were then transmitted at either 3.05 GHz (F1) or 3.75 GHz (F2) to an analogue receiver (TBSI) where they were demultiplexed. The headstage had a 180 mAh battery which lasted about four hours. Accelerometer signals and signal lock indicator were routed to the analogue inputs of the Cerebus Neural Information Processor (Blackrock Microsystems), and neural signals were routed to the front-end amplifier of the Cerebus system. Neural signals were then band pass filtered (0.8 – 7 KHz) and sampled at 30 kHz (.ns6 files). For each channel, a threshold was set at −6.25 below the RMS noise floor. 48 samples around threshold crossings were saved for offline sorting (.nev files). Spikes were sorted using Offline Sorter (Plexon) with the Valley-Seek algorithm to identify units and generate templates, after invalidating noise waveforms and recalculating the principal components. Templates were then used to sort unsorted waveforms. For simplicity, both multi-units and well-isolated single units are reported in the count of neurons found on each array and provide an upward bound for our estimates of array yield for a given recording. SNR for each unit was computed as the peak minus the trough of the mean waveform divided by two times the standard deviation of the waveforms averaged across time. Units with SNR greater than 2.5 were included for each dataset. Recordings used for estimating chronic array yield were performed with the 96-channel Cereplex-m digital headstage (Blackrock microsystems) through the digital hub so that we could sample all 96 channels of the Utah Array. In the case of the second array for marmoset TY, the recordings were performed with the Exilis 96 channel wireless headstage (Blackrock Microsystems).

#### X-ray Reconstruction of Moving Morphology (XROMM)

Two X-ray sources (90 kV, 25 mA) and two image intensifiers (Xcitex ProCapture, 200 fps, 900 x 900) were used to reconstruct the motion of radio-opaque tantalum beads (1 mm diameter, Bal-tec) placed subcutaneously in the marmoset’s torso and upper limb. To implant the tantalum markers, anesthesia was induced with a combination of Ketamine/dexmedatomidine (2 - 6 mg/kg, 75 - 125 µg/kg) and Atipamezole (0.75 - 1.25 mg/kg) to facilitate reversal. Anesthesia was maintained throughout the brief procedure with isoflurane (1 - 3%). We placed markers using angeocatheters (16G, Becton, Dickinson and Company) using the following technique: we introduced the needle subcutaneously, removed the needle while leaving the sheath, placed the marker into the sheath and replaced the needle to expel the marker from the sheath into place under the skin. After the marker was in place, both the needle and sheath were removed carefully to guarantee the marker remained in place. Veterinary adhesive was used to close the angiocatheter entry point when necessary.

We used XMA lab (Knörlein et al., 2016) to track the position of the radio-opaque markers in each camera view, reconstruct the 3D position of each marker in the capture volume, and determine rigid body transformations or endpoint positions for the torso, arm and hand. We describe motion in terms of joint coordinate systems which measure the relative motions of anatomical coordinate systems positioned in the torso, humeral segment of the arm, the forearm, and hand. We defined head orientation as the rotational component of the rigid body transformation computed using singlular value decomposition on the motion of the head markers. We used MAYA (Autodesk) to register these the anatomical coordinate systems to bone models of marmoset head, torso and upper limb.

We determined precision thresholds for estimations of joint angles and marker positions using established methods (Brainerd et al., 2010b; Menegaz et al., 2015), also described by Walker and colleagues (2020). Briefly, a frozen cadaver with a matching marker configuration was moved within the capture volume of the XROMM. Variability in pairwise marker distance and computed joint angles was taken as a measure of error. In a frozen cadaver, inter-marker distances and joint angles should not change. As such any variability should reflect measurement noise. We could track marker position with sub-millimeter precision, and we could record joint angles to within one degree for all measured degrees of freedom, except shoulder rotation for which we could measure to within two degrees.

#### Wireless recordings during unconstrained behavior recordings

To record unconstrained behavior within the entirety of the marmosets’ home cage, we used a webcam (c910, Logitech, 15 fps, 900 x 1200) and an Arduino MEGA to record video of this behavior and synchronize it with neural responses. The Arduino was used to generate a start pulse corresponding to the beginning of the recording and a persisting square wave which was shared with the neural recording system through its analog inputs to facilitate alignment of behavioral video with neural data. Behavioral segmentation was performed manually using ELAN (Brugman and Russel, 2004). Criteria for defining behavioral states are described in Supplementary table 1.

#### Wireless recordings during sleep

For the recordings during sleep, prior to the marmoset falling asleep we replace the fully charged battery and transmitter package. To facilitate this replacement, we preferred doing it when the marmosets were already relaxing in their hammock, so they were relatively stationary. Our second preferred method was to attach the battery by having them stand still with positive reinforcement. Once the battery was in place, we would turn the headstage on briefly to check signal quality and configure the parameters for recording. Once the recording was set up, we would turn off the headstage and leave the marmoset to fall asleep. We would return once the lights had been out for a few hours to turn on the headstage and start the recording. The switch to turn on the headstage is magnetic. As such we could quickly wave a magnetic wand without disturbing the marmoset. Once the recording had started, we would leave to let it run for the life of the battery and return either at the end of the recording or the next morning to remove and replace the battery. Spike sorting was performed as described for the XROMM dataset. Accelerometer signals were smoothed and decimated for display.

#### Deinstrumentation

In the case of an array no longer yielding quality neural signals, we could either replace the array with a new one reusing the same titanium pedestal, or we could remove this pedestal to deinstrument the animal. In one such scenario, under the same anesthetic protocol described above, we made a midline incision and exposed the feet of the pedestal. We did observe some loosening of the bone screws; however, there was significant tissue growth around the feet of the pedestal. After removing all the screws used to secure the pedestal to the parietal bones during the pedestal implantation, this soft tissue growth held the pedestal firmly to the skull and allowed only minimal rotational movement of the pedestal. To remove the pedestal, we first cut the gold wire bundle leading to the array. Then we used a scalpel blade to slowly incise this soft tissue at the juncture of the pedestal feet with the parietal bone. In this way we worked from the posterior most edges of the pedestal feet, under the feet forward, until the pedestal was no longer held to the skull. We then sutured the midline incision and recovered the marmoset from anesthesia. The animal was monitored closely for any signs of pain and distress for two weeks, after which the sutures were removed (Supplementary Figure 5).

### QUANTIFICATION AND STATISTICAL ANALYSIS

#### Kinematics

In this study we report head orientation, shoulder rotation, elbow flexion, and hand position. We reported head orientation as the rotational component of the rigid body transformation resulting from singular value decomposition of the time varying position of fiducial markers in the pedestal assembly. We reported shoulder and elbow motion as the relative movement of the rigid bodies computed for the torso and humerus, and humerus and forearm (treated as a single rigid body), around anatomically defined joint coordinate systems. For instance, elbow flexion was defined as the relative motion of the humerus and forearm around the axis passing through the medial and lateral epicondyle of the humerus.

We estimated precision thresholds for joint angles and marker position using methods described by Brainerd and colleagues (Brainerd et al. 2010; Menegaz et al. 2015), adapted to our setup using a frozen marmoset cadaver with marker placements matching those of the in-vivo study. We recorded trials of the frozen specimen being moved around the XROMM capture volume. We then processed those data with the marker tracking and joint angle estimation pipeline as in the in-vivo study. With the frozen specimen, we expected the distance between markers, the intermarker distances, to remain fixed. Intermarker distance deviations would indicate measurement error. We used the standard deviation of intermarker distances as our marker tracking error metric. We also expected that, as the specimen is frozen, no changes in joint angles should occur. Any joint angular motion observed in the frozen specimen would indicate error accumulated in joint angle estimation. As such, the standard deviation of joint angular motion was our threshold for measurable motion at that joint angle. We estimated the following precision thresholds for joint angles: shoulder abduction: 0.304°; shoulder flexion: 0.208°; long axis rotation: 1.824°; elbow flexion: 0.203°. Soft tissue artifacts were not considered.

#### Unconstrained behavior

Videos of free behavior were manually annotated using ELAN (Brugman and Russel, 2004) to identify behavioral state transitions. We observed 1194 epochs of 14 different behaviors. The six behaviors reported were chosen for further analysis as they comprised more than 90 percent of the recording duration. There were at least 150 examples of sitting, vertical clinging, leaping and locomotion. While there were fewer examples of foraging and food manipulation, epochs of these behaviors tended to be longer in duration and thus provided balanced sampling. To estimate firing rate distributions for each unit, we drew 100 random samples of 500 msec durations of spiking activity from within epochs of each behavior, taking care to avoid samples 500 msec before or after behavioral state transitions. We then computed firing rates for state specific segments of spiking activity using 1 msec bins and a Gaussian kernel with a standard deviation of 50 msec.

#### Sleep

To obtain local field potentials, the broadband 30kHz signal from a single channel was decimated by a factor of 6 and band pass filtered (0.1-1 Hz, 1-4 Hz, 4-7 Hz, 7-15 Hz) using a second order butterworth filter. The spectrogram was computed using the multitaper spectrogram method with a moving window of 0.5 seconds in the Chronux library for MATLAB (Bokil et al., 2010).

#### Evaluating chronic efficacy

Signal-to-noise ratios were computed for each unit as the peak minus the trough voltage of the mean waveform divided by two times the mean standard deviation. The mean standard deviation was defined as follows: the standard deviation of each sample (1 – 48) was computed across all spikes for a given unit. The mean of these 48 standard deviation values was computed as mean standard deviation. Units with signal to noise greater than 2.5 were included.

## ACKNOWLEDGEMENTS

The authors would like to acknowledge the veterinary staff of the University of Chicago who were instrumental to providing care for the marmosets. Additionally, the authors would like to thank Triangle Biosystems for their help adapting the wireless headstage to our needs. The authors would also like the acknowledge the Wisconsin National Primate Center and especially the National Primate Biological Materials Distribution Program for providing opportunities for surgical practice. The authors would also like to acknowledge Callum Ross and the University of Chicago X-ray Reconstruction of Moving Morphology facility. Lastly, the authors would like to acknowledge the paleoCT facility at the University of Chicago, and Courtney Orsbon for CT scanning support.

## AUTHOR CONTRIBUTIONS

J.D.W.: conceived of the work, designed the approach, performed the surgeries, participated in marmoset care and training, collected the data, analyzed the data, wrote and edited the manuscript. F.P.: assisted in surgeries, participated in marmoset care and training, collected the data, edited the manuscript. M.S.: assisted in surgeries, participated in marmoset care and training, collected the data, edited the manuscript. M.N: provided guidance and direction during design of surgical approach, participated in marmoset care, assisted during surgeries, edited the manuscript. J.N.M: conceived of the work, provided guidance during the design of the approach, obtained funding and resources, edited the manuscript. N.G.H.: conceived of the work, provided guidance during the design of the approach, assisted in surgeries, obtained funding and resources, edited the manuscript.

## SUPPLEMENTARY MATERIALS

**Supplementary figure 1.**
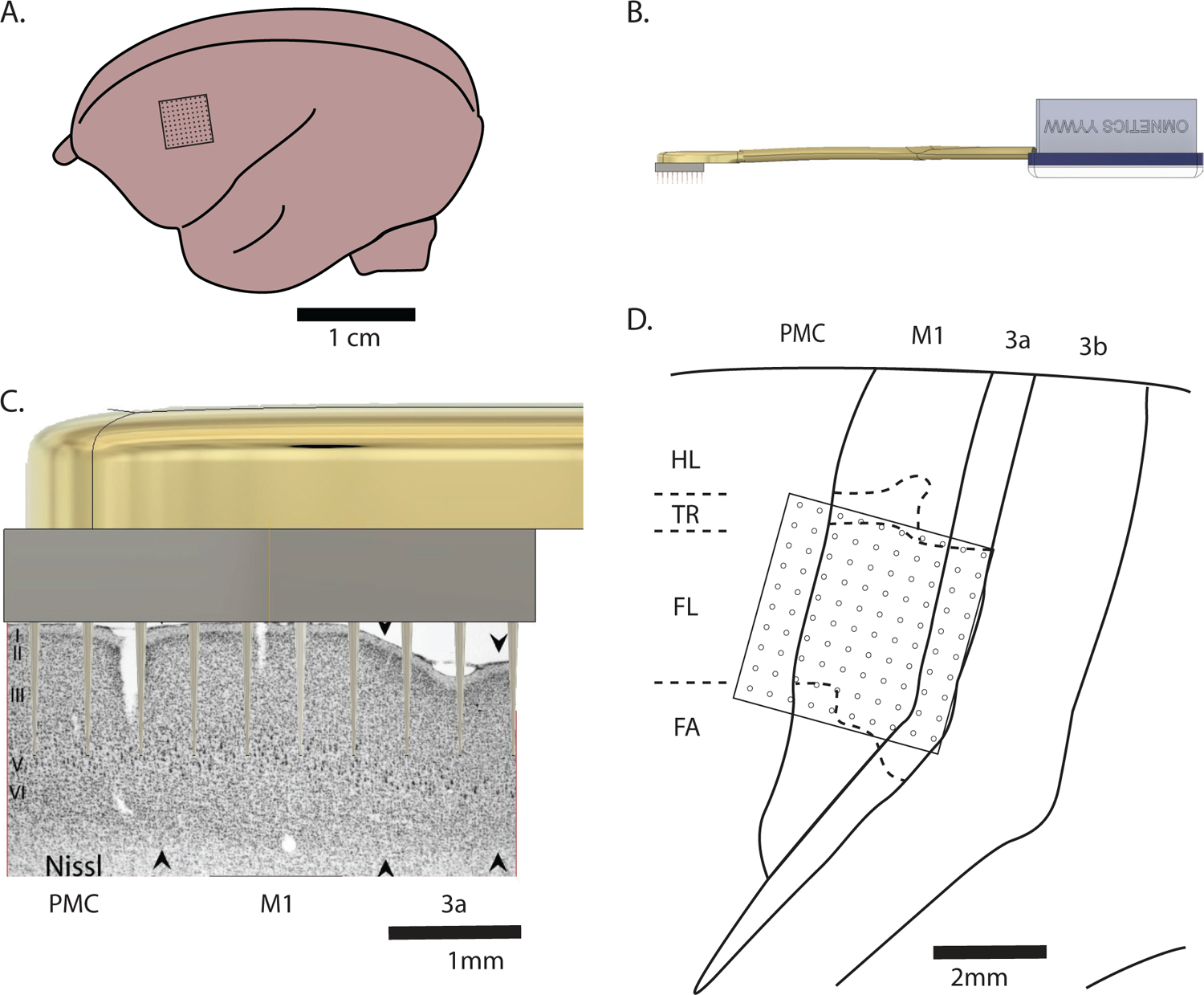
Relevant dimensions of the Utah array and marmoset sensorimotor cortex. A) Illustration of the marmoset brain with footprint of the electrode array targeting forelimb motor cortex (M1). B) Model of the Utah array, wirebundle and Omnetics connector. Scale same as in panel A. C) Utah array with 1 mm electrodes placed within sagittal section of marmoset sensorimotor and premotor cortices. Sagittal section from Burish et al. (2008). D) Outlines of somatotopic organization of marmoset sensorimotor and premotor cortex with footprint of array targeting forelimb M1. Cortical area outlines redrawn from case 6-27 of Burish et al. (2008). Abbreviations: PMC, premotor cortex; M1, primary motor cortex; HL, hind limb; TR, trunk; FL, forelimb; FA, face.

**Supplementary figure 9.**
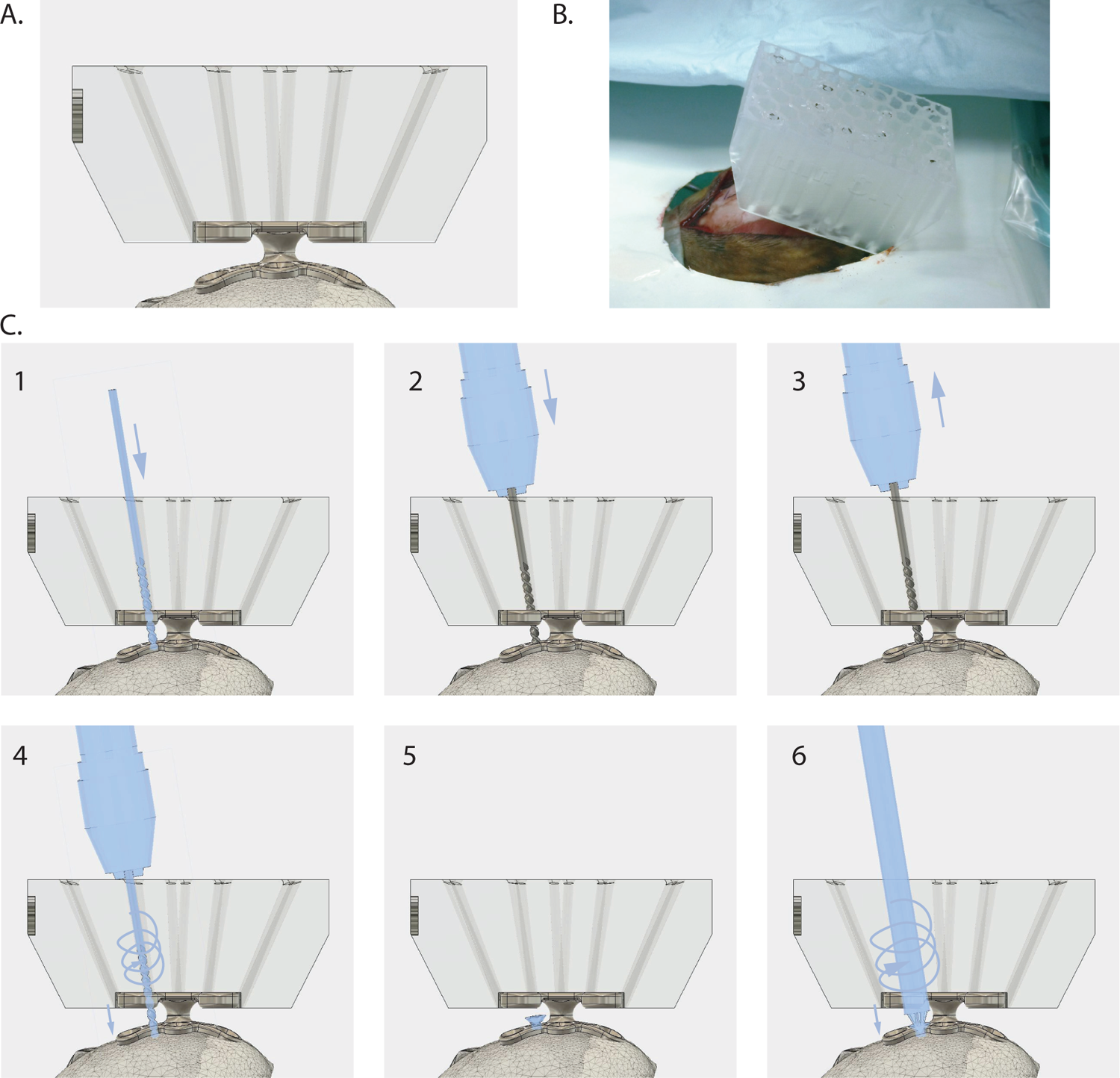
Surgical guide and bone screw placement for pedestal implantation. A) Model of surgical guide mounted on top of the pedestal. Tool paths passing diagonally through the body of the guide allowing alignment of drill and screwdriver with the holes in the feet of the pedestal such that they are perpendicular to the skull at the point of contact. B) Surgical photograph taken after the surgical guide was placed onto the pedestal. C) Sequence of steps used to install each bone screw: 1) Insert drill bit into tool path of guide. 2) Grab bit with pin vise such that the jaws of the pin vice are flush with the top of the guide and the tip of the bit is contacting the bone. 3) Remove the bit from the toolpath with the pin vice. Loosen the jaws of the vice, extend the bit just under 1mm and retighten the jaws of the pin vice. 4) Replace the bit into the same toolpath. While applying constant, but not excessive downward force, rotate pin vice to drill a hole. 5) Place screw above the hole. 6) Insert screwdriver into tool path of the pedestal platform and, while applying constant downward pressure, rotate screwdriver above three turns such that the head of the screw is flush with the dorsal aspect of the pedestal foot.

**Supplementary figure 3.**
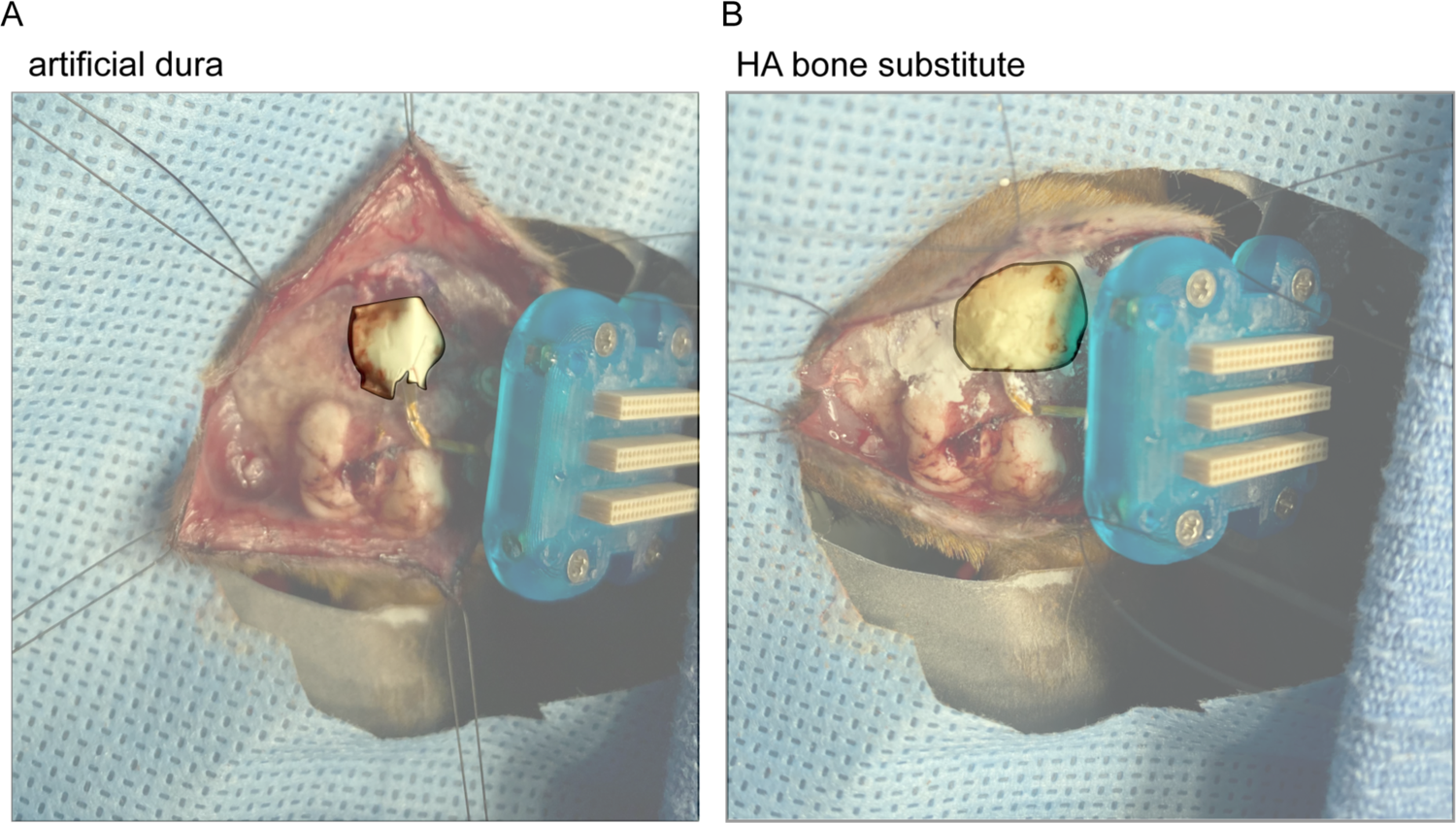
Details of craniotomy closure. A) Photo illustrating use of artificial dura to cover array after insertion in top half of frame. Artificial dura placement was followed by application of either native bone flap and hydroxyapatite bone substitute or as pictured in B) hydroxyapatite bone substitute alone. Note array connector and fixture to the right of the frame and wire bundle medial to craniotomy site, bent toward connector. Also note, in the lower half of the frame, the site of the first array implanted almost four years earlier. The wire bundle for this array has been cut and its connector has been removed, but the array itself was left undisturbed.

**Supplementary figure 4.**
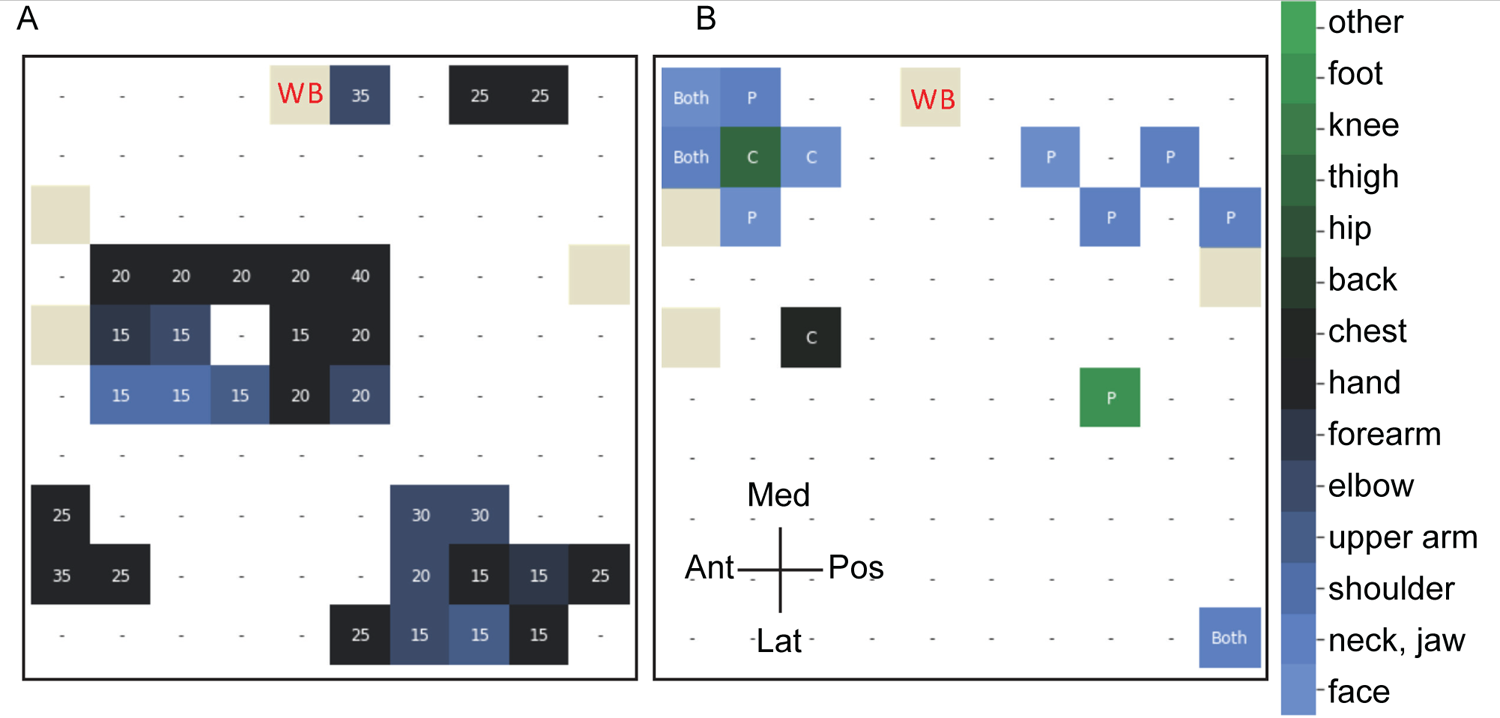
Stimulation effects through electrical, tactile and proprioceptive stimulation are consistent with electrode placement in forelimb M1. A) Map of stimulation effects obtained when stimulating through each electrode on the 10 by 10 grid of electrodes on the array. Each square represents a single electrode. Stimulation effects are described in terms of the primary body part within which electrical stimulation evoked movement. Current threshold for evoking movement effects is given in mA for each electrode. Dashes represent no stimulation effect observed. Tan squares represent reference electrodes. B) Map of neural responses obtained through either tactile stimulation or passive movement of the body parts while the marmoset was lightly anesthetized with isoflurane. Units with high baseline firing rates were not considered, only units that clearly increased their firing rates are reported. Abbreviations: Med, medal; Lat, lateral; Ant, anterior; Pos, posterior; P, proprioceptive; C, cutaneous; WB, wire bundle.

**Supplementary figure 5.**
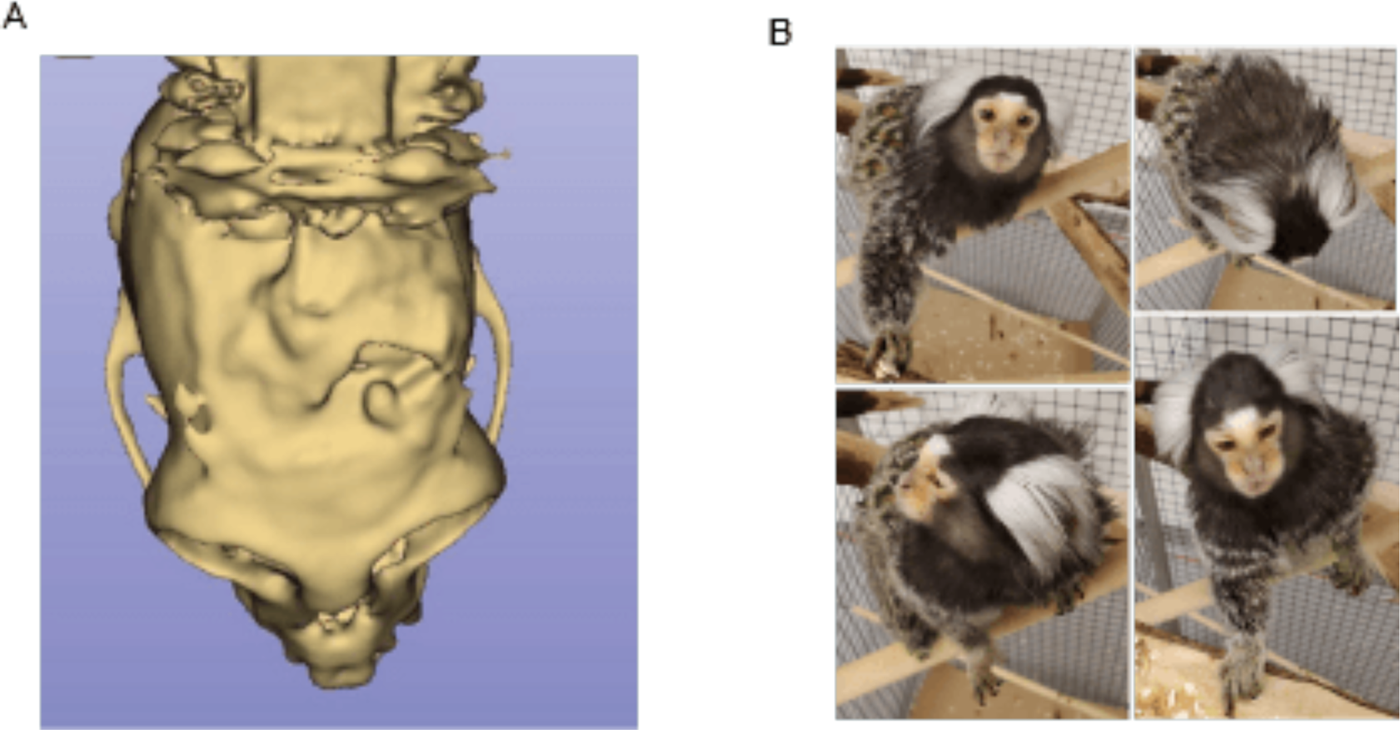
Alternative endpoints: healing and deinstrumentation. (A) Postoperative CT scan showing mostly healed craniotomy. (B) Photos of de-instrumented marmoset after pedestal was explanted, midline incision healed and sutures were removed.

**Supplementary table 1.**
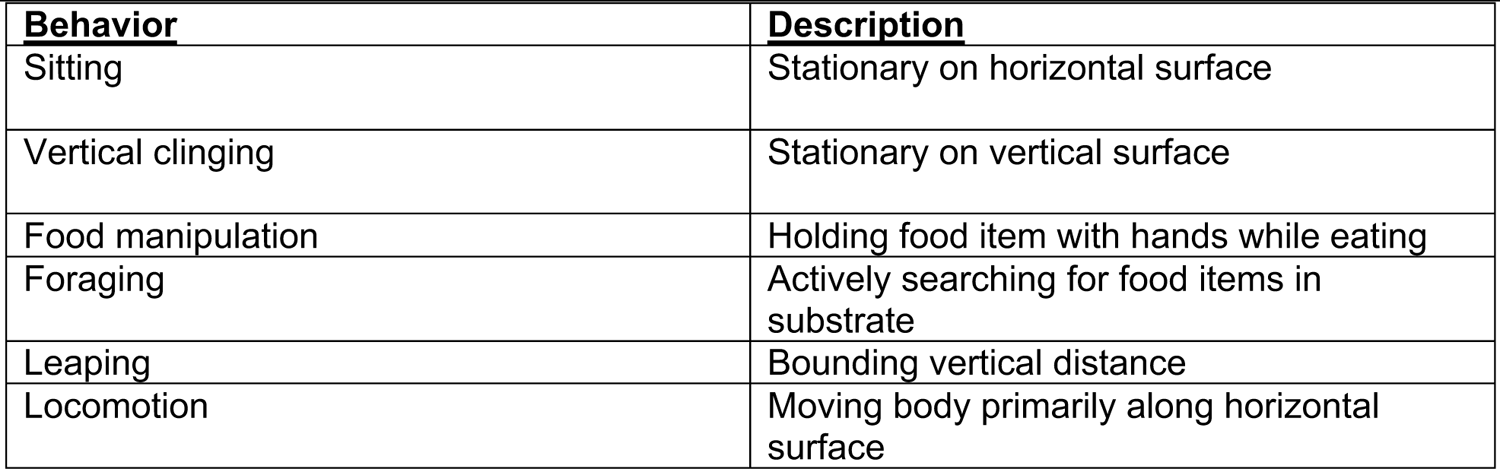
Criteria for behavioral classification. Definition of behaviors used for manual annotation of free behavior analyzed in Figure 5.

| **<u>Behavior</u>** | **<u>Description</u>** |
| --- | --- |
| Sitting | Stationary on horizontal surface |
| Vertical clinging | Stationary on vertical surface |
| Food manipulation | Holding food item with hands while eating |
| Foraging | Actively searching for food items in substrate |
| Leaping | Bounding vertical distance |
| Locomotion | Moving body primarily along horizontal surface |

